# Evolutionary stability of jointly evolving traits in subdivided populations

**DOI:** 10.1101/037887

**Authors:** Charles Mullon, Laurent Keller, Laurent Lehmann

**Affiliations:** Department of Ecology and Evolution, University of Lausanne, 1004 Lausanne, Switzerland

**Keywords:** Social evolution, Infinite-island model, multi-traits phenotypes, evolutionary game theory, behavioural syndromes, dispersal syndromes

## Abstract

The evolutionary stability of quantitative traits depends on whether a population can resist invasion by any mutant. While uninvadability is well understood in well-mixed populations, it is much less so in subdivided populations when multiple traits evolve jointly. Here, we investigate whether a spatially subdivided population at a monomorphic equilibrium for multiple traits can withstand invasion by any mutant, or is subject to diversifying selection. Our model also explores the among traits correlations arising from diversifying selection and how they depend on relatedness due to limited dispersal. We find that selection favours a positive (negative) correlation between two traits, when the selective effects of one trait on relatedness is positively (negatively) correlated to the indirect fitness effects of the other trait. We study the evolution of traits for which this matters: dispersal that decreases relatedness, and helping that has positive indirect fitness effects. We find that when dispersal cost is low and the benefits of helping accelerate faster than its costs, selection leads to the coexistence of mobile defectors and sessile helpers. Otherwise, the population evolves to a monomorphic state with intermediate helping and dispersal. Overall, our results highlight the importance of population subdivision for evolutionary stability and correlations among traits.

## Introduction

One of the main goal of evolutionary biology is to explain patterns of phenotypic diversity, not only between but also within species. Among the most variable and striking phenotypes are social traits, i.e., traits that affect their carriers as well as other individuals in the population, like cooperative or aggressive behaviour. Recent years have witnessed an accumulation of evidence for the co-existence of individuals exhibiting diverse correlated quantitative social traits within populations of insects, mammals, birds, reptiles and fishes (e.g. Sih et al., 2004a; Pruitt et al., 2008; Ducrest et al., 2008; Carter et al., 2010; Cote et al., 2010a; Edenbrow and Croft, 2011; Wang et al., 2013). In sticklebacks, for example, social individuals that tend to be exploratory and aggressive to intruders co-exist with individuals that are less social, but also less exploratory and less aggressive (Laskowski and Bell, 2014). In addition, non-random associations among traits often influence the reproductive success of individuals (reviewed in Sih et al., 2004a; Pruitt et al., 2008), and are to some extent heritable (Drent et al., 2003; van Oers et al., 2004; Sinn et al., 2006; Ariyomo et al., 2013; Wang et al., 2013; Purcell et al., 2014; Dochtermann et al., 2015). These empirical findings have generated considerable interest, yet how selection either leads a population to exhibit little phenotypic variation, or favours phenotypic diversification maintaining correlations among traits, remains in general poorly understood.

The effects of selection on quantitative traits can be studied by looking at the adaptive dynamics of traits, which are the gradual phenotypic changes displayed by a population under the constant influx of mutations (e.g., Eshel, 1983; Parker and Maynard Smith, 1990; Christiansen, 1991; Grafen, 1991; Abrams et al., 1993; Eshel, 1996; Geritz et al., 1998). Unfortunately, a detailed description of adaptive dynamics is in general complicated to achieve. In order to understand the long-term effects of selection on jointly evolving traits, it is therefore more fruitful to study the equilibria of the adaptive dynamics and their evolutionary stability. Importantly, investigating evolutionary stability provides insight into the fundamental features of adaptation in the absence of genetic constraints (Parker and Maynard Smith, 1990), and into the conditions that lead to phenotypic diversification (e.g., random environments, sexual selection, niche partitioning, trophic interactions and social behaviour, van Doorn et al., 2004; de Mazancourt and Dieckmann, 2004; Leimar, 2005; Dercole and Rinaldi, 2008; Brannström et al., 2010).

Evolutionary stability rests on two questions about trait values that are equilibria of adaptive dynamics (also known as singular values, Eshel, 1983; Taylor, 1989; Christiansen, 1991). The first is whether a mutant whose trait is closer to the singular value than the trait of the monomorphic population it arose in will invade. If that is the case, the population will converge to the singular value through recurrent substitutions, and such evolutionary attractors have been coined convergence stable. The second question asks whether a population that is monomorphic for a singular value can resist invasion by any mutant whose trait is close to the singular value. When that is the case, the singular value is said to be locally uninvadable. A singular trait value that is both convergence stable and locally uninvadable is an evolutionary end-point: a population will gradually converge to it and, in the absence of genetic drift or exogenous changes, remain there forever (Eshel, 1983). Alternatively, a convergence stable trait value may also be locally invadable. In that case, the population approaches the convergence stable value but then diversifies, possibly undergoing evolutionary branching, whereby a unimodal phenotypic distribution becomes bimodal, leading to the stable coexistence of highly differentiated morphs (Geritz et al., 1998). Evolutionary branching has been shown to occur in a number of scenarios of social interactions like mutualism, helping or competition (Ferriere et al., 2002; Doebeli et al., 2004; Dercole and Rinaldi, 2008, and references therein). Understanding both evolutionary convergence and uninvadability are therefore necessary to understand how selection leads to phenotypic diversification in social traits.

The evolutionary convergence and local uninvadability of traits evolving in isolation from one another are well-understood in well-mixed populations (or panmixia, e.g., Eshel, 1983; Taylor, 1989; Christiansen, 1991). But to understand patterns of diversification and covariation among multiple traits requires looking at the evolutionary stability of jointly evolving traits. When multiple traits are under selection, selection on one trait may affect selection on other traits (Lande and Arnold, 1983; Phillips and Arnold, 1989; Brodie et al., 1995), and when a population becomes polymorphic for multiple traits, selection can then preferentially target certain combinations of traits over others, thereby creating phenotypic correlations (Phillips and Arnold, 1989). The evolutionary stability of phenotypes that consist of multiple traits is also well-understood in well-mixed populations (e.g., Lessard, 1990; Leimar, 2009). Studies on the uninvadability of multiple traits in well-mixed populations have for instance shown that interactions among traits can facilitate evolutionary phenotypic diversification (Leimar, 2009; Doebeli and Ispolatov, 2010; Svardal et al., 2014; Debarre et al., 2014; Ito and Dieckmann, 2014).

The vast majority of natural populations, however, are spatially-structured. When dispersal is limited, spatial structure creates genetic structure: interacting individuals are more likely to carry identical alleles than individuals sampled at random from the population. As a result, selection depends on the genetic structure and indirect fitness effects (i.e., the effects that traits in neighbouring individuals have on the fitness of a focal individual, Hamilton, 1964; Eshel, 1972; Rousset, 2004). Evolutionary convergence in subdivided populations has been well-studied, and whether a singular trait value is convergent stable can be determined solely from Hamilton’s selection gradient, which uses the probability that two neutral genes are identical-by-descent (i.e., genetic relatedness) as a measure of genetic structure (Frank, 1998; Day, 2001; Rousset, 2004). The joint study of Hamilton’s selection gradients on multiple traits then informs on the convergence stability of multiple traits (Brown and Taylor, 2010, for a review), and this type of analysis has yielded intuitive insights into the evolutionary convergence of many co-evolving social traits, such as dispersal and sex-ratio, altruism and dispersal, altruism and punishment, or altruism and kin-recognition (e.g., Gandon, 1999; Perrin and Mazalov, 2000; Reuter and Keller, 2001; Rousset and Gandon, 2002; Lehmann and Perrin, 2002; Gardner and West, 2004; Leturque and Rousset, 2004).

By contrast, local uninvadability in subdivided populations has received far less attention and is significantly more challenging to study than in well-mixed populations (Day, 2001; Metz and Gyllenberg, 2001; Ajar, 2003; Massol et al., 2009; Wakano and Lehmann, 2014; Svardal et al., 2015). Computational methods that determine uninvadability in subdivided populations are available (Metz and Gyllenberg, 2001; Massol et al., 2009), but they do not straightforwardly reveal how uninvadability depends on genetic structure and fitness effects, which are central components of biological evolution (e.g., Hamilton, 1964; Frank, 1998; Rousset, 2004; Wenseleers et al., 2010, but see Svardal et al., 2015, for environmental effects on uninvadability when local populations are infinitely large). Two studies so far have helped understand the influence of genetic structure and fitness effects by interpreting the uninvadability of a singular trait value in terms of relatedness and indirect fitness effects when local populations are small (Ajar, 2003; Wakano and Lehmann, 2014). In particular, it was shown that whether a trait value is uninvadable depends on how selection on the trait affects genetic structure. However, the works of Ajar (2003) and Wakano and Lehmann (2014) looked at a trait that is evolving in isolation from any other traits. Thus, how population subdivision, which is so important to evolution, influences the diversification and the maintenance of correlations among jointly evolving traits is still poorly understood.

In this paper, we investigate mathematically when multiple evolving traits in a subdivided population are locally uninvadable, or alternatively, when diversification occurs. In the case of diversification, we also study the type of correlations among traits that are favoured by selection. In order to perform our analysis of selection, we use the growth rate of a mutation when rare. We highlight the influence of spatial genetic structure by expressing the conditions for uninvadability, diversification and the among-traits correlations favoured by selection in terms of genetic relatedness coefficients and indirect fitness effects.

## Model

### Life-cycle

We consider a haploid population divided into an infinite number of patches, each with *N* adult individuals (Wright, 1931’s island model). The life cycle is as follows. (1) Patches may go extinct, and do so independently of one another. (2) Each of the N adults in a surviving patch produces offspring (in sufficient numbers for each patch to be always of size N at the beginning of stage (1) of the life cycle), and then either survives or dies. (3) Dispersal and density-dependent competition for vacated breeding spots occur.

This life-cycle allows for one, several, or all adults to die per life-cycle iteration, thereby encompassing overlapping and non-overlapping generations, as well as meta-population processes where whole patches go extinct before reproduction. We assume that each offspring has a non-zero probability of dispersal, so that patches are not completely isolated from each other, and that dispersal occurs to a randomly chosen patch (i.e., there is no isolation-by-distance). However, dispersal is allowed to occur in groups and before or after density-dependent competition, so that more than one offspring from the same natal patch can establish in the same non-natal patch.

### Multidimensional phenotypes

Each individual expresses a genetically determined multidimensional phenotype that consists of *n* continuous traits. These traits can affect any event of the life cycle, under the assumption that the expression of these traits and their effects are independent of age. For instance, the fertility or mortality of an individual is assumed to be independent from its age and that of any other individual who may affect it.

### Uninvadability of a multidimensional phenotype

A resident phenotype **z** = (*z_1_,z_2_, …, z_n_*), where *z_p_* is the value of the *p*^th^ quantitative trait is said to be uninvadable if any mutation that arises as a single copy in the population and causes the expression of phenotype *ζ = (ζ_1_, ζ_2_, …, ζ_n_*), goes extinct with probability one. The uninvadability of a resident phenotype **z** is therefore assessed by considering the evolutionary success of all possible mutations **ζ** that would arise when the population is monomorphic for the resident.

## Lineage fitness, uninvadability and selection on multiple traits

### Lineage fitness and global uninvadability

In order to measure the evolutionary success of a mutation, we use the fact that it is very rare in the population when it arises as a single copy. Because the total number of patches is infinite, it will initially continue to remain rare, even if it increases in frequency due to selection and/or genetic drift and starts to spread to other patches. As a result, interactions among mutants from different patches are very unlikely and can be neglected in the initial growth phase of the gene lineage initiated by the mutation. The evolutionary success of a mutation can therefore be assessed by measuring the individual fitness of a carrier of the mutation that is randomly sampled from the mutantlineage, when the rest of the population is monomorphic for **z**. We define the lineage fitness of a mutant **ζ** which arises as a single copy in a resident **z** population as

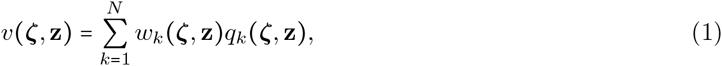

where *w_k_ (ζ, z)* is an individual fitness function that gives the expected number of adult offspring produced by an individual carrying the mutation when there are k mutants in a patch (including the individual itself if generations overlap). The quantity *q_k_(ζ, z)* is the probability that a randomly drawn member of the mutant lineage resides in a patch with a total number *k* of mutants, and which is calculated over the quasi-stationary distribution of mutant patch types in the initial growth phase of the mutant lineage (before extinction or eventual invasion). The lineage fitness of a mutation therefore gives the expected individual fitness of a randomly sampled carrier of the mutation from its lineage (Day, 2001; Lehmann et al., 2015).

If the lineage fitness of a mutation is greater than one, then we expect it to invade the population because in that case, the individual fitness of an average carrier of the mutation is greater than the individual fitness of a resident, which is one. In fact, building on the branching process approach of Wild (2011), we find that *v*(**ζ, z**) provides an exact assessment of the evolutionary success of a mutation because if, and only if *v*(**ζ, z**) ≤ 1, the mutation goes extinct with probability one, but if *v*(**ζ, z**) > 1, there is a nonzero probability that the mutant persists in the population (Appendix A). We can therefore formulate the following uninvadability condition:

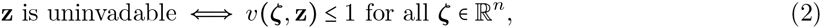

i.e., a phenotype **z** is uninvadable if, and only if, a mutation that causes its bearers to express **z** has the greatest lineage fitness when all other individuals in the population also express **z** (see also, Lehmann et al., 2015).

### Lineage fitness and other measures of fitness

Lineage fitness can be linked to other measures of fitness that have been used to study mutant invasion. The links depend on the properties of the mutant lineage, which are captured by the patch profile distribution, *q_k_*(**ζ, z**).

#### Lineage fitness, average growth rate and basic reproductive number R_0_

In general, *q_k_*(**ζ,z**) can always be expressed as the probability that a member of the mutant lineage, randomly drawn from the quasi-stationary distribution of the lineage, resides in a patch with *k* mutants (i.e., in terms of the eigenvectors of the matrix describing the mean of the branching process before extinction or eventual invasion, Appendix eq. A-8 and Harris, 1963, p.44). In that case, lineage fitness is also equal to the average growth rate of the mutant (Appendix eq. A-2; Caswell, 2001; Tuljapurkar et al., 2003). Condition (2) is then equivalent to the condition that the leading eigenvalue of the linearised deterministic dynamical system describing the growth rate of the mutant when rare in the population is less than one (Caswell, 2001; or less than zero in continuous time, Metz, 2010). In addition, because the basic reproductive number *R*_0_ - the leading eigenvalue of the matrix that gives the total expected offspring production over the local lifetime of the mutant lineage in a patch - is a proxy to the growth rate (i.e., *R*_0_ is sign equivalent to the growth rate around one; Ellner and Rees, 2006), condition (2) is also equivalent to the condition that *R*_0_ ≤ 1 (Appendix eq. A-11).

#### Lineage fitness proxy and R_m_

When a local lineage mutant lineage (i.e., confined to a single patch) can only be initiated by a single founding mutant, it is possible to obtain a proxy for the average growth rate (i.e., an invasion fitness proxy) that is of the same functional form as lineage fitness (Appendix eqs. A-9 - A-21). In that case, the only difference with lineage fitness is that the probability *q_k_*(**ζ,z**) stands for the probability that a randomly drawn member of a local mutant lineage resides in the focal patch when there are *k* mutants (see eq. (20) of the Appendix and Box 2 of Lehmann et al., 2015).

Written as an invasion fitness proxy, lineage fitness is easier to evaluate explicitly because it requires only a matrix inversion rather than the explicit computation of eigenvectors. The expected number *R_m_* of emigrants produced by a local lineage founded by a single immigrant is also an invasion fitness proxy (Appendix eq. A-14; Metz and Gyllenberg, 2001; Massol et al., 2009; Mullon and Lehmann, 2014). Although sign-equivalent to lineage fitness proxy around one, *R_m_* is not equal to lineage fitness proxy. In fact, *R_m_* is equal to *R*_0_ when local lineages can only be founded by a single mutant. Although lineage fitness proxy and *R_m_* can both be used to study uninvadability when local lineages are initiated by a single mutant, lineage fitness proxy is expressed directly in terms of individual fitness *w_k_*, which in some cases make it easier to interpret or manipulate.

### Evolution when mutations have weak phenotypic effects

When mutations have weak phenotypic effects, the lineage fitness of a mutation can be approximated in terms of the deviation between resident and mutant phenotype (**ζ − z**), by way of a Taylor expansion

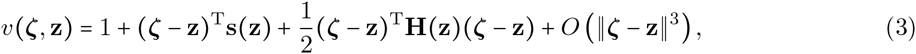

where *O*(||**ζ − z**||^3^) is a remainder of order ||**ζ - z**||^3^. Here, the *n*-dimensional vector **s(z)** is the selection gradient at **z**, i.e., each of its entry *p*,

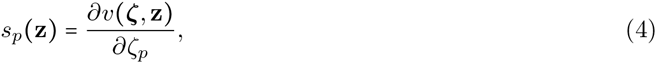

measures the change in lineage fitness due to varying only the trait at position *p*, henceforth referred to as trait *p*, when the population is monomorphic for **z** (all derivatives here and thereafter are evaluated at the resident value **z**). Each (*p, q*) entry of the *n* × *n* Hessian matrix **H(z)**,

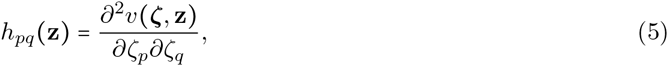

measures how simultaneously varying traits *p* and *q* affect lineage fitness (i.e., the non-additive or interaction effects of *p* and *q* on lineage fitness).

#### Singular phenotype

A multidimensional phenotype **z** is called singular if the selection gradient vanishes at **z**, namely if

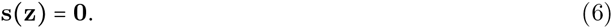

Singularity is a necessary condition for an interior phenotype to be convergent stable and/or uninvadable.

#### Convergence stability

When the difference between a non-singular resident (**s(z) *≠*0**) and the mutant is small (||**ζ - z**|| << 1), the selection gradient in the island model is sufficient to determine whether the mutant will go extinct or fix in the population (Rousset, 2004). When the mutation rate is very low, fixation of a new mutation occurs before another mutant arises - as traditionally assumed in the adaptive dynamics framework (Dercole and Rinaldi, 2008), or the weak-mutation strong-selection regime of population genetics (Gillespie, 1991), and evolution proceeds by a trait substitution sequence whereby the population jumps from one monomorphic state to another. A singular phenotype **z** will then be approached by gradual evolution, i.e., is convergence stable (Leimar, 2009), if the *n* × *n* Jacobian **J(z)** matrix with (*p*, *q*) entry

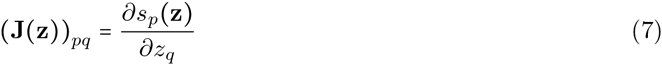

is negative-definite at **z**, or equivalently if all its eigenvalues are negative.

#### Local uninvadability

At a singular resident (**s(z) = 0**), **H(z)** determines whether the resident phenotype is locally uninvadable. From eq. (3), **z** is uninvadable when (**ζ − z**)^T^**H(z)(ζ − z**) < 0 for all **ζ**, i.e., when the matrix **H(z)** is negative-definite, or equivalently when its eigenvalues are negative (e.g., Horn and Johnson, 1985, p. 104). Denoting ⋋_1_(**z**) as the dominant eigenvalue of **H(z)**, we can sum up that **z** is locally uninvadable when

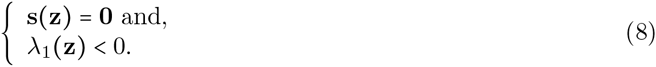

The eigenvalue ⋋_1_(**z**) determines the maximal lineage fitness of a mutant, 1 + ||**ζ−z**||^2^⋋_1_(**z**)/2, at a singular phenotype when the mutant deviates by small magnitude ||**ζ − z**|| from the resident (Appendix B).

#### Diversifying and stabilising selection

Because **H(z)** is symmetric and composed of real entries, if any diagonal entry of **H(z)** is positive, then **H(z)** has at least one positive eigenvalue (Horn and Johnson, 1985, p. 398). It is hence necessary that all the diagonal entries of **H(z)** are non-positive for **z** to be uninvadable. The evolutionary significance of these diagonal entries can be seen by considering the lineage fitness of a mutation that only changes the value of trait *p* by a small amount *δ_p_* = (**ζ − z**)_*p*_. From eq. (3), the lineage fitness of such a mutation at a singular phenotype is 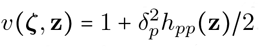. When *h_pp_*(**z**) > 0, selection favours the invasion of any mutation that changes the value of trait *p* (since 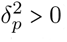), i.e., selection on trait *p* is diversifying or disruptive. Conversely, when *h_pp_*(**z**) < 0, selection on trait p is stabilising. Hence, the sign of the diagonal entries of **H(z)** reflects whether selection on each isolated trait is either diversifying or stabilising (also referred to as concave and convex selection respectively, Phillips and Arnold, 1989).

#### Synergy among traits

As eq. (5) shows, each off-diagonal entry of **H(z)** capture the synergistic effects among pairs of traits on lineage fitness. If *h_pq_*(**z**) is positive, then the effects of trait *p* and trait *q* on lineage fitness are synergistically positive and a joint increase or decrease in both trait values increase lineage fitness. Conversely, if *h_pq_*(**z**) is negative, opposite changes in trait values increase lineage fitness. The mathematical relationship between the eigenvalues of **H(z)** and its off-diagonal entries is less straightforward than with its diagonal entries, but negative eigenvalues, and thus uninvadability, tend to be associated with off-diagonal entries that are close to zero (using results for positive-definite matrices, Horn and Johnson, 1985, p. 398). Therefore, uninvadability is associated with weak synergy among traits (as found in well-mixed populations, Svardal et al., 2014; Debarre et al., 2014).

#### Predicting the build-up of correlations among traits

The synergistic effects among traits on lineage fitness indicate whether selection favours joint or opposite changes in pairs of traits, i.e., when *h_pq_*(**z**) is positive, selection favours a positive correlation among *p* and *q*, and conversely, when *h_pq_*(**z**) is negative, selection favours a negative correlation. This type of selection has thus been referred to as correlational selection (Phillips and Arnold, 1989). From **H(z)**, it is possible to predict how diversifying selection leads to the build-up of phenotypic correlations among *n* traits under scrutiny. As shown in Appendix B, diversifying selection is greatest along the right eigenvector *e*_1_(**z**) that is associated with ⋋_1_(**z**), i.e., the vector such that

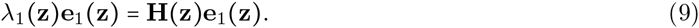

Thus, mutations that are most likely to invade when the resident population expresses a singular phenotype lie on **e**_1_(**z**) in phenotypic space (see Fig. 1 and Appendix B, and as implied by the calculations of Doebeli and Ispolatov, 2010, in well mixed-populations). Thus, the correlations among n traits that are most likely to develop are those given by the direction of **e**_1_(**z**) (Fig. 1).

**Figure 1:**
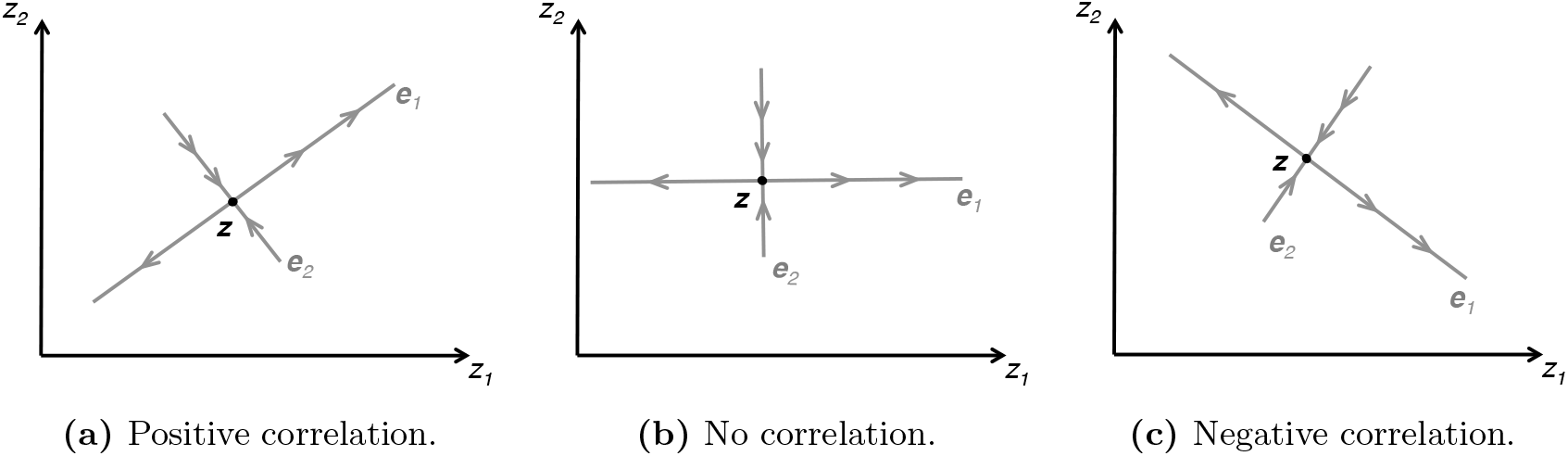
Leading eigenvector and phenotypic correlations favoured by selection. The multi-dimensional phenotype consists of two traits, *z_1_* and *z_2_*. The population is monomorphic for a singular phenotype **z** (black filled circle). The eigenvectors of the Hessian matrix, **e**_1_ and **e**_2_ (grey lines), are positioned to intersect at **z**. A positive eigenvalue, ⋋_1_ > 0, indicates that selection along its associated eigenvector **e**_1_ is diversifying, as shown by the outward arrows. In contrast, a negative eigenvalue, ⋋_2_ < 0, tells us that selection along **e**_2_ is stabilising, as shown by the inward arrows. Selection on phenotypic correlations within individuals depends on the direction of **e**_1_. In (a), the direction of **e**_1_ indicates that selection favours a positive correlation; in (b), no correlation; and in (c), a negative correlation.

The interpretation of **H(z)** in terms of the mode of selection on isolated, and of synergy among, traits mirrors the interpretation of the matrix of second-order effects of selection in well-mixed populations (sometimes denoted *γ* and referred to as the matrix of quadratic selection coefficients, Lande and Arnold, 1983; Phillips and Arnold, 1989; Lessard, 1990; Leimar, 2009; Doebeli and Ispolatov, 2010; Svardal et al., 2014; Debarre et al., 2014). However, it should be noted that in well-mixed populations, the matrix of second-order effects of selection matrix collects the second order effects of traits on individual fitness only (effects on *w* only). By contrast, **H(z)** here summarises the second order effects on lineage fitness, which depend on individual fitness but also on population structure and local demography. In the next section, we highlight how population subdivision affect selection by expressing the second order effects on lineage fitness in terms of individual fitness and relatedness.

## Uninvadability and relatedness

In order to gain greater insight into the effects of population subdivision on selection on jointly evolving traits and uninvadability, as well as connect our results to social evolutionary theory (e.g., Hamilton, 1964; Frank, 1998; Rousset, 2004; Wenseleers et al., 2010), we seek to express the selection gradient and the Hessian matrix in terms of individual fitness and relatedness.

Individual fitness and relatedness can both be recovered from lineage fitness. Lineage fitness depends on *w_k_*(**ζ,z**), which is the individual fitness of a mutant when there are *k* mutants in its patch (eq. 1). More generally, we can write the fitness of mutant and resident alike as *w*(**z_*i*_, z_−*i*_, z**), which is the fitness of individual indexed *i* ∊ {1,&, *N*} in a focal patch with phenotype **z_*i*_**, when the vector of phenotypes among its *N* − 1 neighbours is **z_−*i*_** = (**z_1_,…, z_i−1_, z_i+1_,…, z_*N*_**) and the phenotype carried by all other individuals in the population is the resident phenotype **z**. Then, the fitness of a mutant when there are *k* mutants in its patch that appears in lineage fitness (eq. 1) is

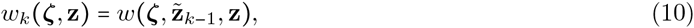

where 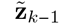 is a vector of multidimensional phenotypes consisting of *k* − 1 entries with phenotype ζ, and *N − k* entries with phenotype **z**. Note that because the population under consideration is not class structured, the fitness of a focal individual is not affected by which precise individual in the patch is a mutant, what matters is how many residents and how many mutants there are in a patch (i.e., 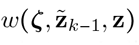 is invariant under permutations of the entries of the vector 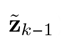). In order to illustrate what an individual fitness function typically looks like, and simultaneously provide a basis for examples to come, we present in Box 1 a fitness function for an iteroparous population.

#### Box I. Individual fitness under a Moran process

A patch goes extinct with probability *e*(**z_*i*_, z_−*i*_, z**). In non-extinct patches, individual *i* produces a large number *f*(**z_*i*_, z_−*i*_, z**) of offspring. Then, exactly one adult individual dies on each non-extinct patch. Individual *i* has a death rate function μ(**z_*i*_, z_−*i*_, z**), so that it dies with probability 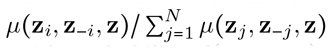 in the current time step. Its offspring disperse independently from one another with probability *d*(**z_*i*_, z_−*i*_, z**), during which they survive with probability *s*(**z_*i*_, z_−*i*_, z**). Then, the fitness of individual *i* is

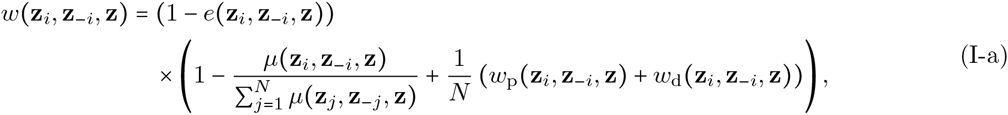

where the term on the first line is the probability that the patch of individual *i* does not go extinct. The first term on the second line is the probability that individual *i* survives to the next generation, while the second term is the expected number of offspring colonising vacant breeding spots, which is decomposed between those that remain in the philopatric patch *w_p_*(**z_*i*_, z_−*i*_, z**)/N, and those that disperse and establish into other patches *w_d_*(**z_*i*_, z_−*i*_, z**)/N. The philopatric component of fitness is given in terms of life-history traits by

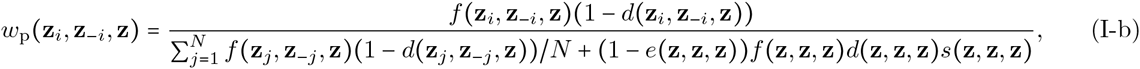

where the numerator is the expected number of offspring of individual *i* that stay in their natal patch and the denominator is the competition faced by an offspring of individual *i* for a philopatric breeding spot. The allopatric component of fitness is given by

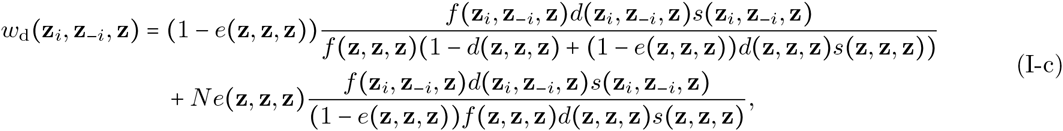

which is the sum between the expected number offspring that colonise non-extinct patches (first line) and those that colonise extinct patches (second line). In non-extinct patches, offspring compete for a single breeding spot with offspring that from that patch, and those that come from other patches. In extinct patches, they compete for *N* breeding spots, which explains the factor *N* in the second summand, and only against offspring that come from other patches.

Lineage fitness also depends on the probability *q_k_*(**ζ−z**) that a randomly sampled member of the mutant lineage has *k* − 1 patch neighbours that are also members of the mutant lineage. The probability mass function *q_k_*(**ζ−z**) characterises identity-by-descent within a patch and therefore relatedness. In fact, the function

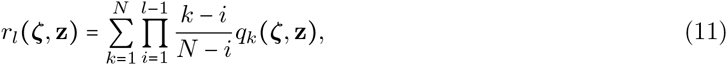

gives the probability that *l* − 1 randomly drawn neighbours without replacement of a randomly sampled mutant from its lineage are also mutants (i.e., that they all descend from the founder of the lineage; for 2 ≤ *l* < *N*). For example, *r*_2_(**ζ,z**) is the probability of sampling a mutant among the neighbours of a random mutant individual, and thus provides a measure of pairwise relatedness between patch members. Under neutrality, all individuals in the patch have the same phenotype (**ζ = z**), and therefore *r_l_*(**z, z**) reduces to the probability of sampling *l* individuals without replacement whose lineages are identical-by-descent, which is the standard *l*^th^ order measures of relatedness for the island model (e.g, Roze and Rousset, 2008, eqs. 22-27). Using the relationships (10)-(11), the selection gradient and Hessian matrix can be expressed in terms of individual fitness and relatedness.

### Uninvadability in subdivided populations

**The selection gradient:** classical kin selection effects. First, we find that the selection gradient on trait *p* can be written as

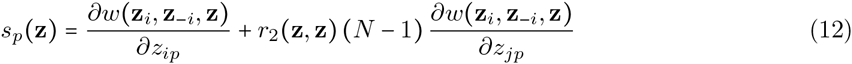

(Appendix C). The first derivative measures the change in fitness of a focal individual as a result of a change in its own trait *p* (i.e., it measures the direct fitness effects of trait *p*). The second derivative measures the change in fitness of the focal due to the change in trait *p* in a patch neighbour (i.e., it measures the indirect fitness effect of trait *p*). Owing to the permutation invariance of patch members on focal fitness (eq. 10), we arbitrarily choose this neighbour to be individual *j ≠ i*. The indirect fitness effect is weighted by the neutral coefficient of relatedness *r*_2_ (**z, z**) between two neighbours (eq. 11). Hence, eq. (12) is the usual selection gradient on a single trait for the island model of dispersal, which is Hamilton (1964)’s selection gradient on trait *p*, or the so-called inclusive fitness effect of trait *p* (Rousset, 2004).

**The Hessian matrix: kin selection effects beyond neutral relatedness.** Next, we find that the (*p, q*) entry of **H(z)** can be decomposed as the sum of two terms,

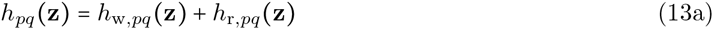

(Appendix D). The first term, *h_w,pq_*(**z**), measures the effect that joint changes of traits *p* and *q* have on individual fitness, while holding the distribution of mutants at neutrality. The second term, *h_r,pq_*(**z**), captures the effect that a change in each trait *p* and *q* has on the local distribution of mutants. As we next see, both terms depend on population subdivision and demography.

The first term of eq. (13a) is

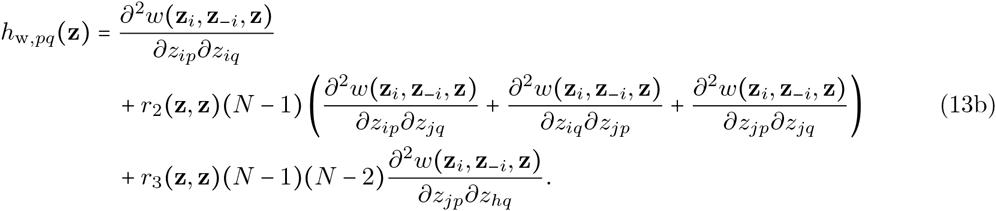

The first derivative measures the change in fitness of a focal individual as a result of a joint change in its traits *p* and *q*. The first and second derivatives on the second line of eq. (13b) measure the change in focal fitness due to a change in one trait of the focal, and a joint change in the other trait of a neighbour. The third derivative on the second line is the change in focal fitness due to a neighbour expressing joint changes in *p* and *q*. Finally, the last derivative of eq. (13b) is the change in focal fitness due to a change in one trait of a neighbour (*j*), and a joint change in the other trait of another neighbour (*h ≠ j*). The fitness effects arising from phenotypic changes in a single neighbour are weighted by *r*_2_(**z, z**), and in two different neighbours by *r*_3_(**z, z**), which is the probability of sampling two mutants without replacement among the neighbours of a random mutant individual (which is also equal to the probability of sampling three mutants without replacement from a random patch under neutrality, **ζ = z**).

Overall, the expression for *h_w,pq_*(**z**) measures the direct and indirect fitness effects due to a joint change in the values of traits *p* and *q* while holding the demography constant. It shows how synergistic effects among two traits can arise due to indirect fitness effects alone: from the fitness effects due to one genetically related neighbour changing both traits, and from those due to two different genetically related neighbours, each changing different traits.

The second term in eq. (13a) is

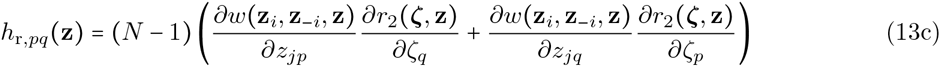

where the first (second) product between derivatives measures the change in fitness to the focal resulting from a patch neighbour changing its trait *p* (*q*), weighted by the change in the probability that a randomly sampled neighbour of a mutant is also a mutant due to a change in trait *q* (*p*).

Eq. (13c) shows that synergy among traits can be due to the combination of indirect fitness effects of one trait, and of a change in mutant relatedness due to a change in another trait value. For example, when a change in *p* in a neighbour causes an increase in a focal mutant’s fitness, and simultaneously a change in *q* increases the probability that the neighbour carry the mutation, eq. (13c) shows that this causes synergy between *p* and *q* to increase, and as a result, so does selection for mutations that change *p* and *q* jointly. Therefore, in subdivided populations, selection can favour a correlation between a trait with indirect fitness effects and another other trait that affect local demography if overall, it results in related individuals reaping greater benefits than unrelated ones.

Eq. (13) for a single trait (*p = q*) substituted into the condition for uninvadability (8) reduces to eq. (B.22) of Lehmann et al. (2015), which was derived under the assumption that only a single mutant can initiate a local lineage. In addition, eq. (13c) for a single trait (*p = q*) is consistent with previous interpretation that the second order effects of selection on an isolated trait depends on how selection affects relatedness (eq. 29 of Wakano and Lehmann, 2014, and under the assumption that generations do not overlap, see second line of eq. 9 in Ajar, 2003). Hence, our analysis not only confirms previous conditions for uninvadability of a single trait, but also shows that these hold under more general conditions, allowing for instance local patch extinctions or dispersal in groups from the same patch. But more importantly, we have extended previous analyses to consider selection on multiple traits under limited dispersal. This highlights that interactions among traits, which are important to uninvadability and selection on phenotypic correlations, can arise through the effects of traits on genetic structure and on the fitness of neighbours. We now proceed to study how uninvadability is computed explicitly, in particular showing how the effect of selection on relatedness is evaluated (i.e., *∂r_2_(ζ, z)/∂ζ_p_*).

## The Moran process and weak selection

### Uninvadability under a Moran process

If the population follows a Birth-Death Moran process, individual fitness is as in Box 1. Since offspring disperse independently from one another in this model, local lineages can only ever be initiated by a single individual in a patch (see below eq. A-12 in Appendix A). We can therefore use lineage fitness proxy (**“Lineage fitness proxy and R_m_”** section). For simplicity, we also assume in this section that there is no patch extinction. Pairwise and three-way neutral relatedness are found using standard techniques (e.g., Karlin, 1968) and are given in Table 1. Together with the fitness function (eq. I-a in Box 1), they allow for the evaluation of the selection gradient *s_p_*(**z**) (eq. 12), and for the *h_w,pq_*(**z**) (eq. 13b) component of the Hessian matrix.

**Table 1:** Backward dispersal, neutral relatedness and spatial scale of competition for a Moran life-cycle in the absence of patch extinction. *m*(**z**) refers to the backward probability of dispersal (i.e., the probability that a breeding spot is filled by a dispersing offspring in a monomorphic population Gandon, 1999), which may depend on the evolving phenotype **z**. ^†^ See Appendix F for derivation. The coefficient *a*(**z**) is the spatial scale of competition when the evolving trait affect payoffs, which in turn affect fertility. It is found by Taylor expanding individual fitness to the first order of ∊, and re-arranging to take the form of Appendix eq. (G-1). If payoffs affect other life history traits, like adult survival, then *a*(**z**) will take a different form.

|  |  |
| --- | --- |
| $m(\mathbf{z})$ | $\frac{s(\mathbf{z}, \mathbf{z}, \mathbf{z})d(\mathbf{z}, \mathbf{z}, \mathbf{z})}{1 - d(\mathbf{z}, \mathbf{z}, \mathbf{z}) + s(\mathbf{z}, \mathbf{z}, \mathbf{z})d(\mathbf{z}, \mathbf{z}, \mathbf{z})}$ |
| $r_2(\mathbf{z}, \mathbf{z})^\dagger$ | $\frac{1 - m(\mathbf{z})}{1 + m(\mathbf{z})(N - 1)}$ |
| $r_3(\mathbf{z}, \mathbf{z})^\dagger$ | $\frac{2(1 - m(\mathbf{z}))^2}{(2 + m(\mathbf{z})(N - 2))(1 + m(\mathbf{z})(N - 1))}$ |
| $a(\mathbf{z})$ | $\frac{(N - 1)(1 - m(\mathbf{z}))^2}{N - (1 - m(\mathbf{z}))^2}$ |

The remaining term necessary for the second order effects of selection, *h_r, pq_*(**z**) (eq. 13c), depends on the first order effect of trait *p* on pairwise relatedness, which we find is

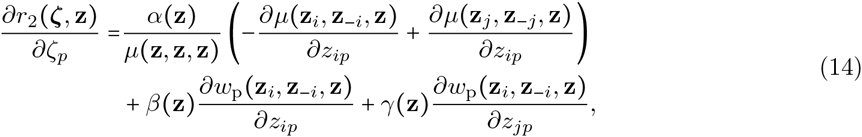

where *μ*(**z_*i*_,z_−*i*_, z**) is the death rate of individual *i* and philopatric fitness *w_p_*(**z_*i*_,z_−*i*_,z**) is its expected number of offspring that remain in the natal patch (Box 1, Appendix E for derivation of eq. 14). The functions α(**z**), β(**z**) and γ(**z**) are positive and decrease with the neutral measures of relatedness (expressed in terms of demographic parameters in Table 2). The second line of eq. (14) shows that mutant relatedness increases if the trait causes the mutant lineage to locally grow faster than a resident lineage, consistent with the effect on relatedness in a Wright Fisher model (Appendix eq. E-23 for link between eq. 14 and Wright-Fisher model). Because generations overlap in the Moran model, relatedness among mutants is different than among residents when the mutant affects the chance of individuals to survive from one generation to the next and this is captured by the first line of eq. (14). This first line shows that relatedness increases if a change in a trait decreases the death rate of its carrier, but increases that of a randomly sampled patch neighbour. Such a trait results in the longer coexistence of multiple mutant generations in the same patch, and therefore increases relatedness between mutants.

**Table 2:** Weights for the first order effects of traits on relatedness for a Moran life-cycle in the absence of patch extinction (eq. 14). See Appendix G for derivation.

|  |  |
| --- | --- |
| $\alpha(\mathbf{z})$ | $\frac{N(N - 1)m(\mathbf{z})(1 - m(\mathbf{z}))}{(2 + m(\mathbf{z})(N - 2))(1 + m(\mathbf{z})(N - 1))^2}$ |
| $\beta(\mathbf{z})$ | $\frac{N}{(1 + m(\mathbf{z})(N - 1))^2}$ |
| $\gamma(\mathbf{z})$ | $\frac{2N(N - 1)(1 - m(\mathbf{z}))}{(2 + m(\mathbf{z})(N - 2))(1 + m(\mathbf{z})(N - 1))^2}$ |

Eqs. (12)-(13) and (14) provide all the necessary components to characterise the uninvadability of multi-dimensional phenotypes in subdivided populations under a Moran life-cycle. We note that if the evolving traits affect only adult fertility and/or offspring survival, philopatric fitness can be written as

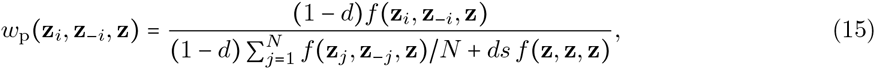

where *f*(**z_*i*_, z_−*i*_, z**) is the number of offspring produced by individual *i*, *d* is the probability that an offspring disperses and s is the probability that it survives dispersal (Box 1). Substituting eq. (15) into eq. (14) shows that when **z** is a singular phenotype, traits have no effects on relatedness, i.e.,

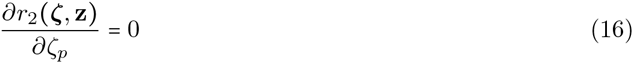

(Appendix eq. E-25). So, when assessing the uninvadability of **z** under a Moran process, the effect of traits on relatedness can be ignored if there are only fecundity effects, thereby facilitating mathematical analysis. As we show in the next section, uninvadability conditions can also be made simpler for a large class of demographic models when traits have weak effects.

### Uninvadability under weak selection

Within an arbitrary life cycle as laid out in the Model section, suppose that as a result of social interactions within patches, individuals receive a material payoff, like food shelter, immunity from pathogens, or some material resources. The expected material payoff obtained by individual *i* in a focal patch during social interactions is written π(**z_*i*_, z_−*i*_, z**) if its phenotype is **z**_*i*_, its *N* − 1 neighbours have phenotypes **z**_−*i*_, and the remainder of the population has resident phenotype **z**. We assume that the evolving traits only affect the payoffs received during social interactions within patches, and that the payoffs affect only weakly life history traits, such as fecundity, adult survival or dispersal.

A given life history trait, written as g(z_*i*_, z__*i*_, z) for a focal individual with phenotype z_*i*_, can then be expressed as

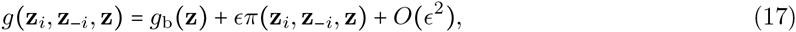

where *g*_b_(**z**) is a baseline value for all individuals that may depend on the resident population, and ∊ > 0 is the effect of payoff on the life history traits, which is assumed to be small. We assume that the fitness *w*(**z_*i*_, z_*−i*_,z**) of the focal individual increases with it material payoff, but decreases or is unaffected by the material payoffs of its neighbours. Also, it is assumed that the effect on the fitness of the focal of changing the payoffs of a single of its neighbour is weaker than the effect of changing the payoffs of the focal.

When the above assumptions holds, individual fitness can be expressed as a linear function of ∊ (Appendix eq. G-1). As a consequence, we find that the entries of the selection gradient s(**z**) are

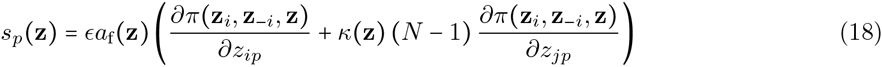

(where *a_f_*(**z**) is positive and model-dependent, see Appendix eqs. G-1 - G-6). This expression closely resembles the general selection gradient eq. (12), with the first term within brackets capturing direct fitness effects, and the second, indirect effects. However, instead of being expressed in terms of the individual fitness function (*w*(**z_*i*_, z_*−i*_, z**)), eq. (18) depends directly on the payoff function, and instead of *r*_2_(**z**), the indirect effects of selection are weighted by the quantity *κ*(**z**), which is a scaled measure of relatedness among two individuals that balances the effects of relatedness and local competition (e.g., Queller, 1994; Lehmann and Rousset, 2010; Akcay and Van Cleve, 2012; Van Cleve, 2015). Relatedness and local competition tend to have opposite effects on selection on social traits as the former promotes the evolution of traits with positive indirect fitness effects whilst the latter favours the evolution of traits with negative indirect fitness effects. Scaled relatedness is therefore useful towards understanding how the balance between relatedness and local competition affect social evolution (Queller, 1994; Lehmann and Rousset, 2010). The explicit expression and interpretation of *κ*(**z**) are given in Box 2 (eq. II-b).

##### Box II. Scaled relatedness and competition

Each scaled relatedness term, *k(**z**), ι(**z**),* and η(**z**) balances relatedness with local competition as,

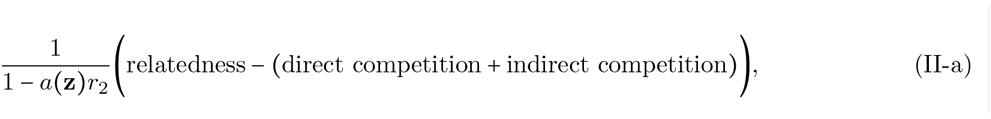

where 0 ≤ *a*(**z**) ≤ 1 measures how much of the competition is between individuals of the same patch. When *a*(**z**) is small, competition is mostly global, when *a*(**z**) is close to 1, competition is mostly local. It is found according to the fitness function (Appendix G, eq. G-1). Scaled relatedness therefore discounts the effects of local competition from the effect of relatedness on selection. In

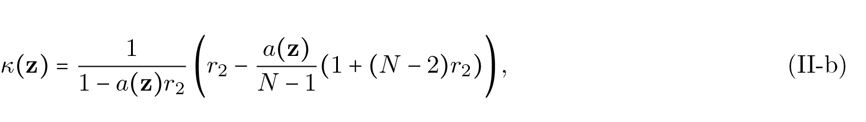

the factor 1 within 1 + (*N − 2)r_2_* captures direct competition among two individuals within a patch because when the focal interacts with an individual *j*, it directly affects its payoff, and thereby directly affects the competition for its own offspring. E.g., if **z** is an altruistic trait, ∂π(**z_i_, z_−*i*_, z**)/∂*z*_*jp*_ is positive, so the focal increases the fertility of *j* and increases the local competition for its own offspring. The term (*N − 2)r_2_* reflects that the neighbour *j* also affects the payoff of the other (*N* − 2) individuals of the patch, and if those are related to the focal (with probability *r*_2_), this indirectly affects competition among related mutants. The term

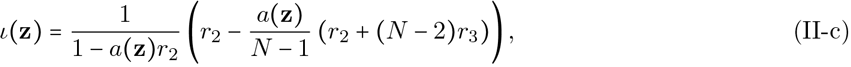

is a scaled measure of relatedness term among two individuals that incorporates the local competition arising from the interactions among two mutants. The term *r*_2_ within *r*_2_ + (*N − 2)r_3_* reflects the competition arising from the effects of the focal on the payoffs of *j* when they are related. The second term within the bracket reflects that *j* also changes the payoffs of the other individuals in the patch related to itself, and that in turn this affects the competition for the focal if they are all related to the focal, which occurs with probability *r*_3_. Similarly, in

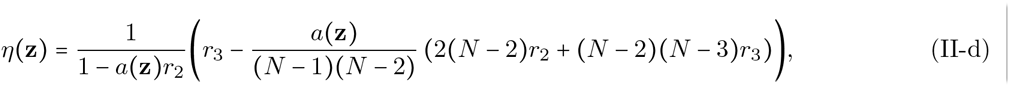

which is a scaled measure of relatedness term among three individuals, the term 2(*N − 2)r_2_* is the effect of the competition that emerges from the focal interacting with individuals, and that interaction affects the payoffs of a third individual. Finally, the term (*N − 2)(N − 3)r_s_* is the indirect competition among related mutants that arises as consequence of neighbours related to the focal interacting with two other individuals related to the focal.

Importantly, we find that when traits have weak effects on fitness, they have no effect on pairwise relatedness, *∂r_2_*(**ζ, z**)/∂ζ_*p*_ = 0. As a result, the second component of the Hessian matrix vanishes (*h_r,pq_*(**z**) = 0), and the (*p, q*) entry of **H(z)** is

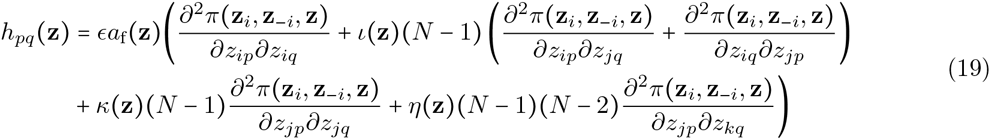

(Appendix eqs. G-7 - G-8). This equation bears close resemblance to eq. (13b), and its elements can be interpreted similarly. However, eq. (19) depends directly on the payoff instead of fitness functions, and on two additional scaled relatedness measures, which, like *κ(**z**)*, balance the effects of relatedness and local competition: ι(**z**), between two randomly sampled individuals; and η(**z**) between three individuals (see Box 2 for a detailed interpretation of these coefficients; and when there is only one evolving trait, *n* = 1, and interactions are pairwise the term in parentheses in eq. 19 reduces to eq. 37 of Wakano and Lehmann, 2014).

Eqs. (18)-(19), along with Box 2, is all that is necessary to evaluate the uninvadability of multidimensional phenotypes in subdivided populations, provided traits only affect the payoffs received during interactions, and selection is weak. Because eqs. (18)-(19) depend on the payoff rather than the fitness function, they tend to be easier to explore mathematically. In addition, the expressions for scaled relatedness reveal more clearly the effects of demography on two antagonistic forces on social traits: relatedness and competition among kin.

However, one should be cautious that the equilibrium values found for life history traits using eq. (17) may not correspond to the equilibrium values that would be found by considering the evolution of the life history trait itself. For instance, while it is possible to model the evolution of game strategies when payoffs affect the ability to disperse using eqs. (17)-(19), equilibrium strategies found in this case may not predict the same equilibrium values for dispersal if we modelled the evolution of dispersal itself. When directly modelling the evolution of dispersal, it is not possible to use the simpler eqs. (18)-(19) because dispersal typically affects the probability that individuals of the same patch carry the same mutation (i.e., *∂r*_2_(**ζ, z**)/∂ζ_*p*_ ≠ 0). We show this explicitly in the next section by considering the evolution of dispersal itself, together with another social trait.

## The joint evolution of helping and dispersal

In order to illustrate the above results and how to apply them, we now study the joint evolution of helping and dispersal. The evolutionary paths of helping and dispersal are intimately intertwined (Lehmann and Perrin, 2002; Le Galliard et al., 2005; Hochberg et al., 2008; El Mouden and Gardner, 2008; Purcell et al., 2012; Parvinen, 2013) because in subdivided populations, the level of dispersal determines relatedness, and therefore tunes selection on helping traits (e.g., Rousset, 2004). Simultaneously, dispersal evolution directly responds to the level of kin competition within a patch (Hamilton and May, 1977).

The complicated interaction between helping and dispersal has meant that so far, theoretical studies of the evolution of these two traits have either focused on evolutionary convergence and ignored the problem of uninvadability (Lehmann and Perrin, 2002; Le Galliard et al., 2005; Hochberg et al., 2008; El Mouden and Gardner, 2008), or relied on simulations and numerical methods to study invadability (Purcell et al., 2012; Parvinen, 2013). Using our framework, we are able to analytically study not only the joint invadability of helping and dispersal, but also the correlation among those two traits that are favoured by selection when evolutionary branching occurs.

### Biological scenario

The life cycle is assumed to follow the Moran process in the absence of patch extinction (see section Moran process and Box 1). During the adult stage, individuals pair up randomly and engage with one another in the so called “continuous snow drift game”, which is a model of helping behaviour that can lead to evolutionary branching among helpers and defectors (Doebeli et al., 2004; Wakano and Lehmann, 2014). In this model, the payoff received by individual *i* when it interacts with *j* is expressed as

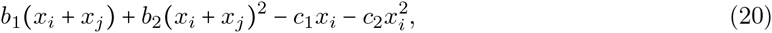

where 0 ≤ *x*_*i*_ ≤ 1 is the level of helping expressed by individual *i*, and *x_j_* that of individual *j*. The constants *b*_1_ and *b*_2_ tune the benefit to *i* of both interacting partners investing into helping, while *c*_1_ and *c*_2_ tune the cost to *i*. Individual fertility increases with the average payoff received,

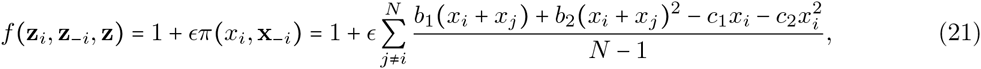

where the parameter ∊ *>* 0 measures the effect of payoffs on fertility. After reproduction, an adult in each patch is selected at random to die, with each adult having the same death rate. The offspring of an individual *i* disperses with probability *d_i_ ∊* (0,1]. Dispersing offspring survive during dispersal with probability *s*.

### Set-up

From these assumptions, the fitness of individual *i* is given by Box 1, with no patch extinction (e(**z_*i*_, z_*−i*_,z**) = 0), constant adults survival (μ(**z_*i*_, z_*−i*_,z**) = μ), and constant survival during dispersal (*s*(**z,z,z**) = *s*). The vector of trait values of an individual *i* consists of its level of helping and dispersal probability, **z**_i_ = (*x_i_, d_i_*). The vector of phenotypes in the rest of the patch is **z**_*−i*_ = ((*x*_1_, *d*_1_),…, (*x*_*i*−1_,*d*_*i−i*_), (*x*_*i*+1_,*d*_i+1_),…, (*x_N_, d_N_*)), and **z** = (*x, d*) stands for resident strategies. Hence, trait *p* = 1 corresponds to helping and trait *p =* 2 corresponds to dispersal. The neutral relatedness functions, *r*_2_(**z,z**) and *r*_3_(z,z), are those given in Table 1. We now have all the elements to use our framework, and start by considering the evolution of each trait in isolation, and then study their coevolution.

### Uninvadability of helping with fixed dispersal

Substituting fitness (eq. I-a, Box 1) into the selection gradient (eq. 12) for helping, we find that when dispersal is fixed at d for all individuals, there is a unique singular helping strategy for which the selection gradient vanishes, which is

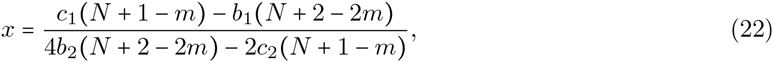

where *m* = *m*(**z**) is the probability that a breeding spot is filled by a dispersing offspring (i.e., the backward dispersal probability, see Table 1). The invadability of this singular point is assessed by calculating *h*_11_(**z**), which is found by substituting fitness (eq. I-a, Box 1) into eq. (13) and evaluating it at eq. (22). Although such a computation yields an analytical expression for *h*_11_(**z**), it is too complicated to generate useful insights. We therefore first study the invadability of helping under weak selection, i.e., with e small in eq. (21), and using eq. (19) to compute *h*_11_(**z**) at eq. (22). Using the spatial scale of competition given in Table 1, and the scaled relatedness coefficients of Box 2, fitness (eq. I-a, Box 1) into eq. (19) gives

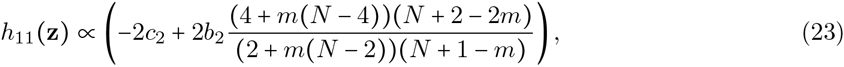

where we have ignored e and a_f_(z) since they are both positive and we are only interested into whether *h*_11_(**z**) < 0 or not. Eq. (19) shows that the singular point (eq. 22) is uninvadable, i.e., *h*_11_ < 0, only if

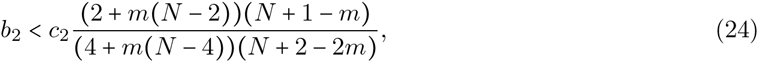

where the factor of *c*_2_ on the right hand side is positive and decreases with backward dispersal (*m*) and patch size (*N*), so that it correlates positively with patch relatedness.

When dispersal is complete (*d* = 1, so that *m* = 1), eq. (24) reduces to the results previously found for well-mixed populations: the singular helping strategy (eq. 22) is uninvadable only if *b*_2_ < *c*_2_ (Doebeli et al., 2004). In other words, the benefits of helping should accelerate at the same rate or slower than its cost. Otherwise, the accelerating returns of helping favour the diversification of helping strategies, and leads to the evolutionary branching between helpers and defectors (Fig. 3, top left panel).

As dispersal decreases (*d* < 1, so that *m* < 1), relatedness among individuals within patches increases, and indirect effects become increasingly important in the fate of newly arising mutations, consequently affecting the stable level of helping. Insights into the effects of population subdivision on the stability of helping can be obtained by expressing *h*_11_(**z**) close to full dispersal (m ∽ 1),

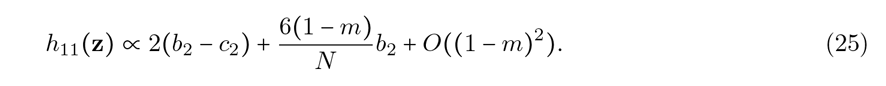

The first and second summands of eq. (25) respectively capture direct and indirect fitness effects. The latter increases with relatedness, here measured by (1 - *m)/N*. Indirect effects also increases with *b*_2_ because the payoffs to a focal individual increase by a factor of *b*_2_ whenever neighbours change their helping strategy. So if *b*_2_ is positive, indirect effects favour the invasion of mutations that change the degree of helping, and thereby destabilise the singular helping strategy. Conversely, if b2 is negative, then focal payoffs decrease, and indirect effects disfavour any change in helping strategy, stabilising the existing degree of helping.

For a given patch size, eq. (24) shows that there is a threshold value for dispersal under which helping, otherwise invadable in well-mixed populations, becomes uninvadable. For example, when *N* = 4, if *m* < 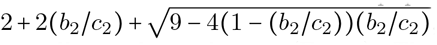, helping is uninvadable in subdivided populations but invadable in well-mixed populations (Fig. 2). This is consistent with previous results found for the Wright-Fisher process (Wakano and Lehmann, 2014), and highlights that when the benefits of helping are decelerating (*b*_2_ < 0), indirect fitness effects combined with high relatedness promote the stability of helping in the population. Because relatedness is greater under the Moran model, the threshold value for dispersal is greater than under the Wright-Fisher model.

**Figure 2:**
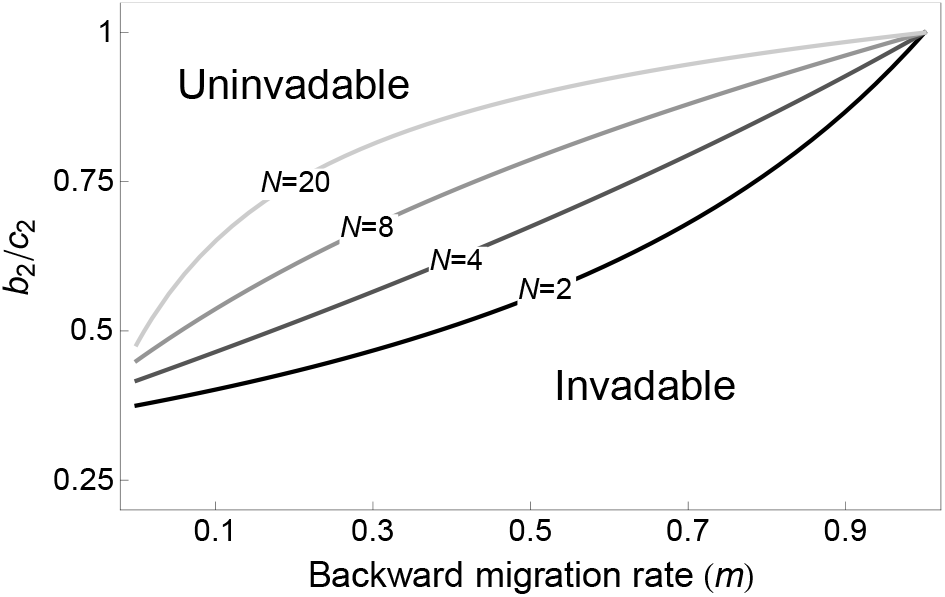
Uninvadability of helping in subdivided populations. Region above the curves gives combination of values of *b*_2_/*c*_2_ and backward migration rate *m* for which helping is uninvadable, while region under the curve gives values for which helping is invadable. Different curves correspond to different patch sizes. Thus, small patch size and low migration rate stabilise helping in subdivided populations.

**Figure 3:**
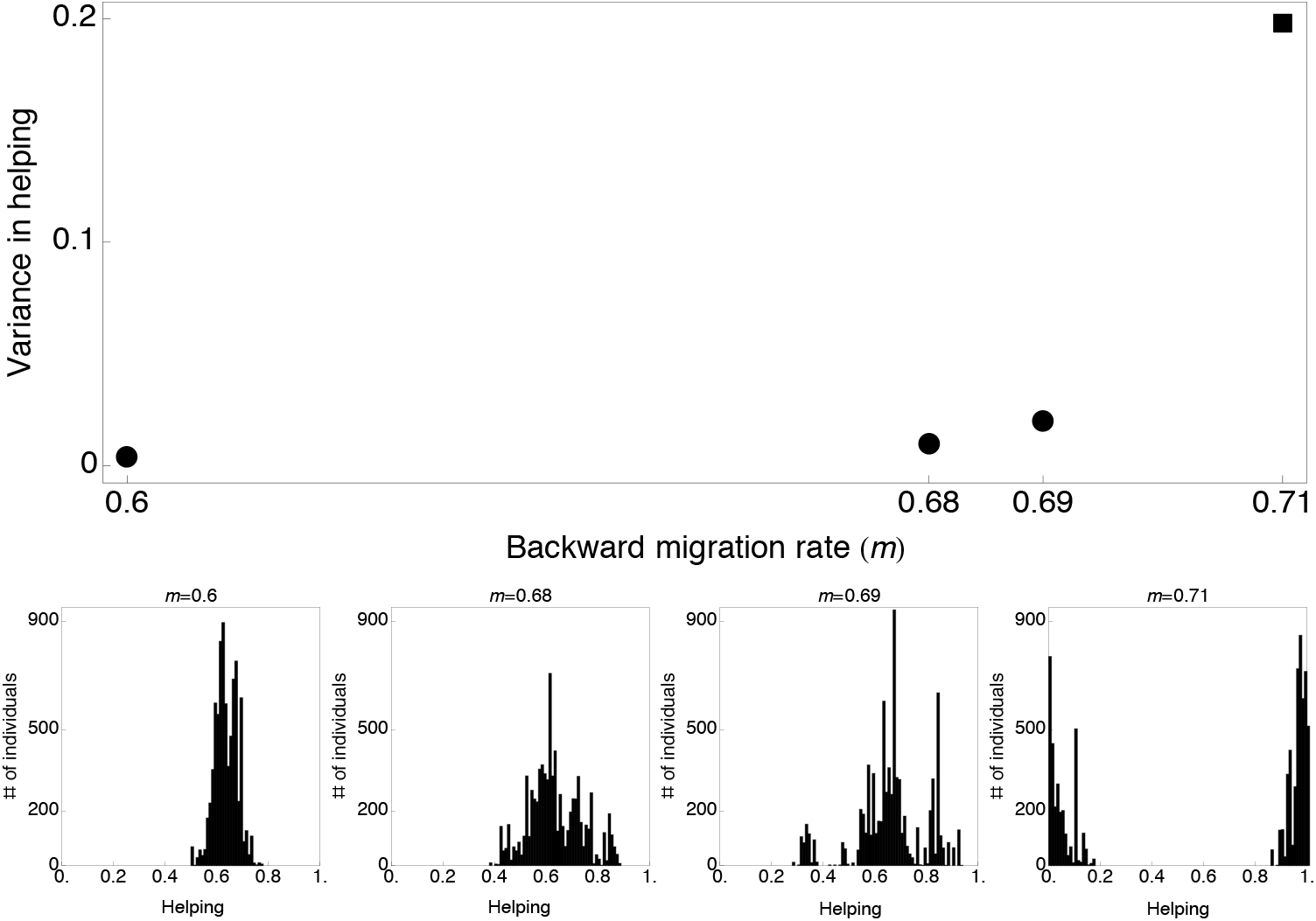
Evolution of helping in a subdivided population. The top panel shows the phenotypic variance of helping in a simulated population for different values of *m* (other parameters were held at *N =* 8, *b*_1_ = 6, *b*_2_ = −1-4, *c*_1_ = 4.56, *c*_2_ = −1.6, number of patches = 1000, Appendix I for details on simulations). The variance is averaged over 5 x 10^3^ generations after 1.45 x 10^5^ generations of evolution. Circles indicate variance for parameter values under which *h*_11_ (eq. 23) is negative, predicting a stable monomorphic population, while squares correspond to a positive *h*_11_, predicting a polymorphic population. The bottom panel shows snapshots of the the level of helping in a simulated population after 1.5 x 10^5^ generations for four different values of *m*.

In order to check the robustness of our weak selection conclusions against increased selection strength, we generated random values for the model parameters, and asked whether the sign of *h*_11_ given by eq. (23) were of the same the sign as *h*_11_ calculated under strong selection (∊ = 1). Both expressions showed the same sign for all combinations of values tested (100% of 465943 trials, Appendix H), suggesting that the sign of *h*_11_ given by eq. (23) is an excellent predictor of the sign of *h*_11_ under strong selection. In addition, individual based simulations with strong selection behave as predicted by the analysis performed under weak selection (Fig. 3). This suggests that uninvadability condition computed under weak selection for traits that affect fertility is robust to increases in selection strength.

### Uninvadability of dispersal with fixed helping

Substituting fitness (eq. I-a, Box 1) into the selection gradient eq. (12) with its derivatives with respect to the probability of dispersal, we find a single singular dispersal strategy,

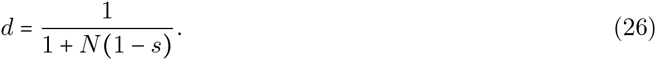

Eq. (26) does not depend on helping because all individuals exhibit the same level of helping and there is thus no variation in offspring production among patches. Since kin competition decreases with patch size (*N*), the candidate dispersal strategy decreases with *N* and since the cost to dispersal decreases with the survival rate during dispersal (*s*), the candidate dispersal strategy increases with *s*. Qualitatively, eq. (26) is therefore the same as the one obtained by the classical models of dispersal evolution, which assume a Wright-Fisher reproductive process (Frank, 1998; Gandon and Rousset, 1999). However, because there is more competition among kin when generations overlap, the singular dispersal strategy under the Moran model is always greater than under the Wright-Fisher model.

Computing *h*_22_(**z**) from eq. (13) and evaluating it at eq. (26), we find that the singular strategy (eq. 26) is uninvadable only if

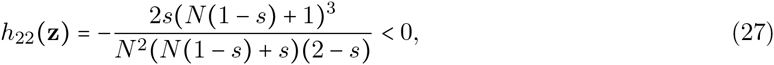

which is always true. Therefore, when dispersal is the only evolving trait, dispersal is always stable in a Moran population. This corroborates the results found under the Wright-Fisher process (Ajar, 2003), but the result for the Moran process is more straightforward, with eq. (27) simpler than the Wright-Fisher condition (eq. 15 in Ajar, 2003). Interestingly, in a Moran population monomorphic for the uninvadable dispersal level, pairwise relatedness is simply

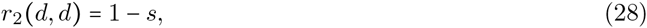

the probability of surviving dispersal.

### The coevolution of helping and dispersal

When helping and dispersal evolve jointly, singular strategies are found by solving simultaneously for vanishing selection gradients for both traits, *s*_1_(**z**) = 0 and *s*_2_(**z**) = 0, which produces the singular point

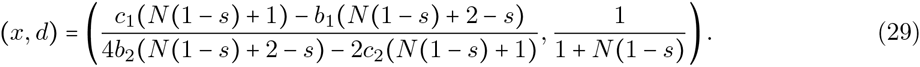

The singular dispersal strategy is the same as in eq. (26), while the singular helping strategy is eq. (22) with *m* = *m*(**z**) as in Table 1, and dispersal d given by the singular phenotype eq. (26).

The joint stability of helping and dispersal is deduced from the leading eigenvalue of **H(z)** (eq. 13), which is a complicated function of the model parameters. However, when the effects of helping on fertility are weak (∊ small in eq. 21), it is possible to express the eigenvalues of **H(z)** as a perturbation of the simpler eigenvalues of **H(z)** with ∊ = 0 (see Appendix J). Using this method, we find that the eigenvalues of **H(z)** under weak selection on helping are

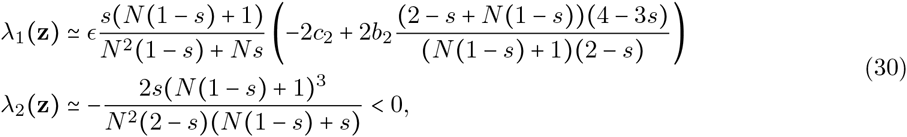

which, respectively, are proportional to eq. (23) and *h*_22_ (eq. 27) at the singular dispersal strategy (i.e., with *m* and *d* as in Table 1 and eq. 29 respectively). Therefore, when the effects of helping on fertility are weak, the condition for helping and dispersal to be jointly stable is the same as the condition for helping to be stable when dispersal is held fixed at the singular dispersal strategy (eq. 24).

Numerical comparisons between the sign of ⋋_1_ in eq. (30) and the sign of the exact leading eigenvalue of **H(z)** calculated under strong selection (with ∊ = 1) shows that the former is a very good predictor of the latter, with both having the same sign in 99.86% (of 464686 cases, Appendix H), and therefore suggests that the uninvadability condition (24) should also hold when the effects of helping on fertility are strong. In addition, individual based simulations with strong selection (∊ = 1) match theoretical predictions very well (Fig. 4).

**Figure 4:**
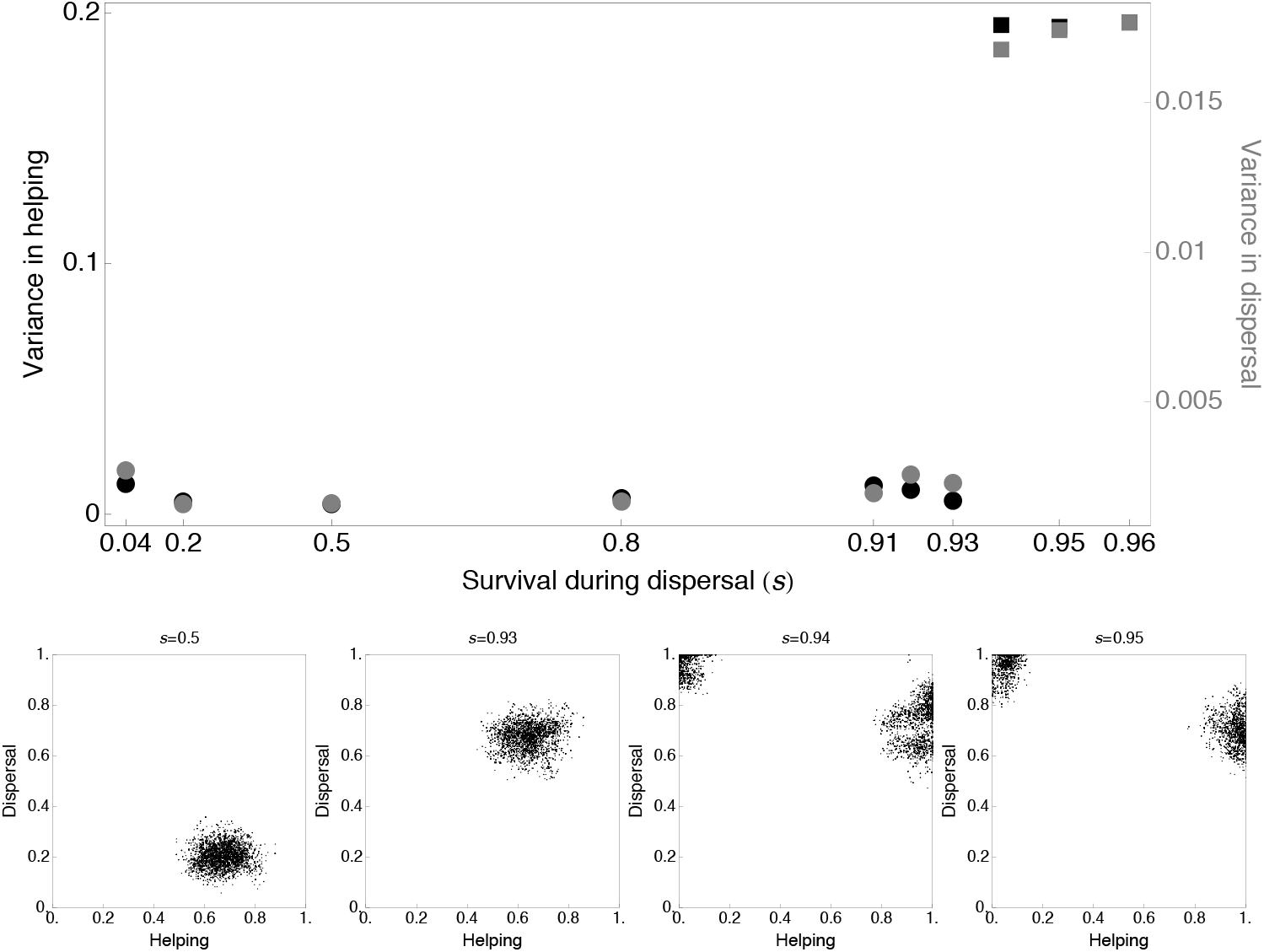
Coevolution of helping and dispersal in a subdivided population. The top panel shows the phenotypic variance of helping (black, left scale) and dispersal (grey, right scale) in a simulated population for different values of *s* (other parameters are as in Fig 3, Appendix I for details on simulations). The variances are averaged over 5 x 10^3^ generations after 1.45 x 10^5^ generations of evolution. Circles indicate variance for parameter values under which the leading eigenvalue of the Hessian matrix is negative, predicting a stable monomorphic population, while squares correspond to a positive eigenvalue, predicting a polymorphic population. The bottom panel shows snapshots of the phenotypes in a simulated population after 1.5 x 10^5^ generations for four different values of s. Each point represents the phenotypic values in helping and dispersal of an individual.

Our results therefore show that invadability of helping strategy also leads to invadability in dispersal strategy. In order to predict whether helpers and defectors evolve different dispersal strategies, we now study the level of correlation among helping and dispersal that is favoured by selection at the singular phenotype. Recall that among-traits correlation are given by the direction of the eigenvector associated with the leading positive eigenvalue of **H(z)** (section **“Predicting the build-up of correlations among traits”**). Since ⋋_2_(**z**) (eq. 30) is always negative, ⋋_1_(**z**) is the leading eigenvalue. If the effects of helping are weak (small ∊), then it is possible to express the eigenvector associated with ⋋_1_(**z**) as a perturbation of the eigenvectors of **H(z)** in the absence of helping (∊ = 0, Appendix J). We find that in this case, the leading eigenvector has direction

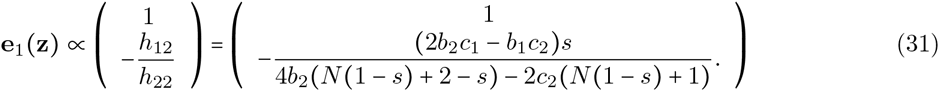

Eq. (31) shows that, since *h*_22_ < 0, selection promotes a positive (negative) correlation among helping and dispersal if the synergy *h*_12_ among them is positive (negative). In order to garner greater insight into the synergy *h*_12_, we approximate it as

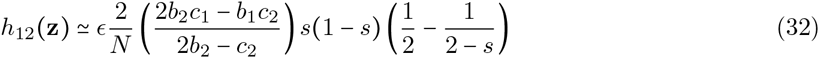

to the first order of ∊, and neglecting terms of order *O*(1/*N*^2^) (Appendix eq. H-1). Looking at eq. (32), we see that the term inside the first parentheses is the effect that a change in the degree of helping of neighbours has on focal payoff at the singular level of helping: (*N* − 1)∂π(*x*_i_, *X*_−i_)/*∂x_j_* = (2*b*_2_*c*_1_ - *b*_1_*c*_2_)/(2*b*_2_ - *c*_2_) + *O*(1/*N*); i.e., the indirect effects on focal’s payoff. Since helping always has positive indirect effects, this term is positive. Then, because the rest of eq. (32) is negative, synergy among helping and dispersal on fitness is always negative.

Since synergy among helping and dispersal is negative, we expect that if polymorphism arises in the population, selection will favour a negative correlation among helping and dispersal. A closer look at the synergy reveals why this is so. Eq. (32) is negative due to the large negative term -1 /(2 - *s*), which stems from the fact that an increase in dispersal has a negative effect on relatedness,

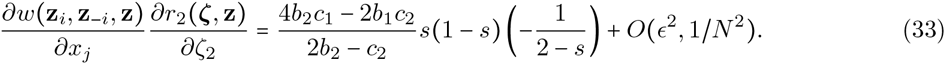

Therefore, selection promotes a negative correlation among helping and dispersal because a simultaneous increase in helping and dispersal leads to greater benefits to unrelated individuals, and, conversely, a simultaneous decrease in helping and dispersal leads to lesser benefits to related individuals. By contrast, lesser dispersal coupled with greater helping leads to greater benefits to related individuals. We find by numerical simulations that the sign of eq. (32) is a very good predictor of the sign of the exact *h*_12_ (**z**) value with arbitrary population size and under strong selection (∊ = 1, Appendix G), so that we also expect a negative correlation among helping and dispersal under strong selection. In fact, individual based simulations under strong selection show that when evolutionary branching occurs, the population splits into very mobile defectors and more sessile helpers (Fig. 4), thereby befitting our analytical predictions.

Our analytical finding of a negative correlation among helping and dispersal corroborates previous simulation results when patches do not vary in size (Purcell et al., 2012; Parvinen, 2013). When patches experience different demographies, simulations show that it is possible for selection to favour a positive correlation among helping and dispersal because helpers and defectors experience different benefits from dispersing: helpers benefit from invading patches with few individuals whereas defectors benefit from invading patches with a large number of individuals (Parvinen, 2013). Other works that have looked at the correlation among helping and dispersal have done so by studying either the convergence stable level of helping according to dispersal strategy (El Mouden and Gardner, 2008), or the convergence stable level of dispersal according to helping strategy (Hochberg et al., 2008), and found results similar to the ones mentioned here. However, it should be noted that in those studies, the presence of helpers and defectors or of disperser and non-dispersers was a starting point and a built-in assumption, rather than a product of evolution like in our analysis.

## Discussion

Understanding uninvadability is key to understand the evolution of quantitative traits because uninvad-ability determines whether selection favours a population to remain monomorphic or become polymorphic (e.g., Eshel, 1983; Taylor, 1989; Christiansen, 1991; Geritz et al., 1998). So far, how genetic structure and indirect fitness effects influence uninvadability had only been studied when single isolated traits evolve (Ajar, 2003; Wakano and Lehmann, 2014), but organisms consist of a multitude of traits that rarely, if ever, evolve in isolation from one another (Lande and Arnold, 1983; Phillips and Arnold, 1989). Here, we have presented a framework to study the uninvadability of phenotypes that consist of multiple quantitative traits, as well as the among traits correlations arising from diversifying selection, in subdivided populations when local populations can be of any size and are connected by limited dispersal.

In order to analyse the effects of selection under limited dispersal, we used the lineage fitness of a mutation, which is the expected number of mutant copies in the offspring generation that are produced by an individual randomly drawn from the mutant lineage (see Lehmann et al., 2015, Akcay and Van Cleve, 2016 and Lehmann et al., 2016 for other applications of this fitness concept). Lineage fitness thus gives an average individual fitness over the possible genetic backgrounds in which a carrier of the mutation can reside. It allowed us to reveal the effects of genetic structure through relatedness coefficients and of indirect fitness effects on the uninvadability of multiple traits. We found that relatedness and indirect fitness effects influence the synergy among traits (or strength of correlational selection). Because strong synergy tends to disfavour uninvadability and thus promote diversification, relatedness and indirect fitness effects may be critical for the evolution of multiple traits under limited dispersal. In particular, we showed that synergy among two traits can arise through the effects of one trait on genetic structure and the indirect fitness effects of the other trait. When positively correlated change in two traits results in related individuals reaping greater fitness benefits than unrelated ones, synergy tends to be positive, and conversely, when negatively correlated changes in two traits result in related individuals reaping greater fitness benefits, synergy tends to be negative.

We further found that, since the synergy among pairs of traits determines the among-traits correlations that develop when polymorphism arises, relatedness and indirect fitness effects also influence the evolution of phenotypic correlations. Because behavioural traits tend to have the greatest indirect fitness effects, our results suggest that synergy due to the combination of traits’ effects on genetic structure and indirect fitness effects is important to the evolution of behavioural syndromes (correlations among behavioural traits within individuals, Sih et al., 2004a, b). Previous studies have suggested that behavioural syndromes may be maintained mechanistically by pleiotropic mutations (Ducrest et al., 2008), and by fitness tradeoffs between life-history traits (Wolf et al., 2007; Reale et al., 2010). Here, our model shows that selection promotes a positive correlation among two traits when one trait has positive indirect benefits, and the other trait increases pairwise relatedness (i.e., when two individuals that show an increase in the value of the trait have a greater probability of being related than two resident individuals).

In this context, dispersal syndromes, which refer to patterns of covariation between the tendency to disperse and other traits, are of particular interest (Clobert et al., 2009). Dispersal syndromes are ecologically and evolutionarily relevant as they influence the demographic and genetic consequences of movement (Edelaar and Bolnick, 2012; Jean Clobert et al., 2012). In Western bluebirds, for example, individuals that disperse further away from their natal site also tend to be more aggressive towards conspecifics and towards sister species, and this has caused a shift in the range of these two species (Duckworth and Badyaev, 2007; Duckworth and Kruuk, 2009). Since dispersal decreases relatedness, our model predicts that the tendency to disperse should be negatively correlated with behaviours that have positive indirect fitness effects. In our model of helping, we indeed found that when evolutionary branching occurs, the population splits between full defectors that always disperse and full helpers that are more sessile. This negative correlation among helping and dispersal also matched previous simulation results when patches do not vary in size (Purcell et al., 2012; Parvinen, 2013). Similarly, our model predicts dispersal to be positively correlated with behaviours that have negative indirect fitness effects, like aggressiveness. Interestingly, both a negative correlation between dispersal and pro-social behaviour (Ims and Ims, 1990; Mehlman et al., 1995; O’Riain et al., 1996; Sinervo and Clobert, 2003) and a positive correlation between dispersal and aggressive behaviour have been observed in natural populations of voles, mole rats, rhesus macaques, mosquitofish and side-blotched lizards (Myers and Krebs, 1971; Mehlman et al., 1995; Cote et al., 2010b; Aguillon and Duckworth, 2015, see also Cote et al., 2010a for review).

Our work also paves the way for more in-depth understanding of uninvadability and evolutionary branching in subdivided populations. A difficulty in studying uninvadability in subdivided population stems from the necessity to calculate the effects that selection on traits has on pairwise relatedness. In this endeavour, our work is helpful in three ways. First, we have explicitly calculated the effects of selection on relatedness for the Moran process, therefore opening the door to studying uninvadability under this standard model of reproduction. Second, we have shown that the leading effects of selection on relatedness are zero at a singular phenotype when the evolving traits only affect adult fertility or offspring survival before dispersal under the Moran process, thus facilitating stability analysis for this case. Third, and more generally, the effect of selection on relatedness has no bearings on uninvadability when traits influence material payoffs that in turn weakly affect fitness (i.e., weak selection), and this holds for any demographical model that fits our general assumptions (see “Main assumptions” section). As illustrated by our study of the joint evolution of helping and dispersal, this method is useful to reach analytical results. In addition, we were able to relate the uninvadability of traits directly to the effects of traits on payoff and to the level of local competition when fitness effects are weak. Since relatedness and local competition have antagonistic effects on the evolution of social traits, our decomposition is useful to understand the forces at play for the uninvadability of social traits. We expect that our method to study uninvadability under weak selection should be particularly useful to understand adaptation in multiple strategies within complex, multi-strategies, games in the presence of genetic structure. Despite our progress, calculating the effect of selection on pairwise relatedness for arbitrary selection and demographical models under our general assumptions remains challenging, and explicitly characterising uninvadability may require extensive additionalcomputations depending on model choice.

In order to reach tractable results, we have made a number of assumptions. Many of them - infinite population size, clonal reproduction of haploid genomes, rare mutation with weak effects - are common to those of the adaptive dynamics framework, and have been extensively discussed elsewhere (e.g., Geritz and Kisdi, 2000; Geritz and Gyllenberg, 2005; Champagnat et al., 2006; Dercole and Rinaldi, 2008). We have also assumed that the population is subdivided according to Wright (1931)’s classic infinite island model. This model ignores important biological realities like isolation-by-distance or temporal and spatial heterogeneity in environment. By changing genetic structure in space, isolation-by-distance makes selection on social traits more complicated (Rousset, 2004). Yet, most relevant insights on how selection moulds social traits are revealed by the infinite island model (like for cooperation and altruism, Lehmann and Rousset, 2010), and we therefore expect results qualitatively similar to ours under isolation-by-distance, at least when patches are homogeneous. It would be interesting to extend our framework to study the effects temporal and spatial heterogeneity in environment on the uninvadability of traits, in particular when traits can themselves influence the environment locally (Rousset and Ronce, 2004). As shown by simulations, temporal heterogeneity in patch sizes due to dispersal patterns may be important for the synergy among helping and dispersal (Parvinen, 2013). More generally, in order to study age-, sex-or caste-specific effects, or demographic stochasticity, it is necessary to incorporate class-structure (i.e., when the effect of a trait on the fitness of an individual is contingent on the class the individual belongs to). Like the vast majority of stability analyses of adaptive dynamics, our model falls short of predicting the course of evolution once evolutionary branching has occurred. This would require describing the post-branching phenotypic distribution in the population (as in Sasaki and Dieckmann, 2011).

In conclusion, we have provided a framework to study the evolutionary stability of intertwined traits in subdivided populations. In particular, with uninvadability expressed in terms of relatedness coefficients and indirect fitness effects, our model sheds light onto the consequences of population structure for evolutionary stability, diversification and the evolution of correlations among traits.

## Appendix

### A. Lineage fitness and uninvadability

We show here that a mutation causing the expression of phenotype ζ that initially appears as a single copy in a resident population expressing **z** will vanish with probability one if, and only if, the lineage fitness of this mutation is less or equal to one (*v*(**ζ, z**) ≤ 1, eq. 1).

#### Multitype branching process

We denote by *X_i_(t)* the random number of patches in the population with *i ∊ I = {1,2, …, N}* mutants at time *t*, which are collected into the vector *X(t) = (X_1_(t),…, X_N_(t))*. We assume that the stochastic process *{X(t)}_t≥o_* is a multi-type branching process (as in Wild, 2011), which is equivalent to assuming that when the mutant is globally rare, only residents can immigrate to a patch that already contains mutants. Under this assumption, the expected change in *X(t)* over one generation is governed by a matrix **A(ζ, z)**, called the mean matrix, whose *(i,j)* element, denoted *a_ij_***(ζ, z**), gives the expected number of patches with *i ∊ I* mutants that are generated by a patch when it has *j ∊ I* mutants, and the population is otherwise monomorphic for **z**,

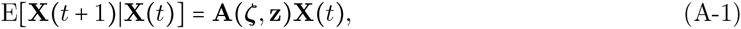

where the expectation is taken over replicates of the one-generational demographic process (i.e., over one life cycle iteration).

It follows from standard results in stochastic demography that the average growth rate of the mutant population is the leading eigenvalue of **A(ζ, z)** (Caswell, 2001; Tuljapurkar et al., 2003). More precisely, the leading eigenvalue of **A(ζ,z)**, denoted *r*(**A(ζ, z)**), gives the time-averaged mean cumulative growth over different replicates or sample paths of the invasion dynamics, i.e.,

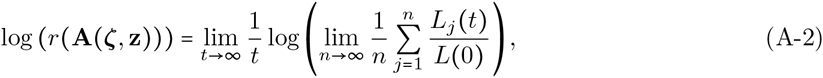

where *L_j_ (t)* is the total random number of patches with at least one mutant at generation *t* for replicate population *j* (i.e., the sum of the *X_i_(t)*’s for a given replicate *j*), *n* is the total number of replicate populations, and *L*(0) = 1 when the mutation arises as a single copy. Since the expected number of patches of each type grows asymptotically with *r*(**A(ζ, z)**), the expected total mutant lineage size, 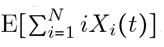, also grows asymptotically at rate *r*(**A(ζ,z)**). Furthermore, the mutant lineage goes extinct at some time *t* > 0 with probability one (i.e., Pr{X(*t*) = 0 for some *t* |**X**(0) = (1,0,…, 0)} = 1) if, and only if the leading eigenvalue of **A(ζ, z)** is less or equal to one (Karlin and Taylor, 1975; Harris, 1963, p. 41), i.e., if, and only if *r*(**A(ζ, z)**) ≤ 1.

#### Lineage fitness as the average growth rate of the mutation

We now show how the growth rate *r*(**A(ζ, z)**) can be expressed as lineage fitness (eq. 1). Writing **u(ζ, z)** as the right eigenvector of **A(ζ, z)** associated with *r*(**A(ζ, z)**), we have by definition

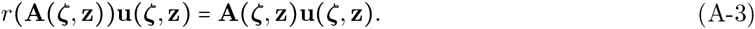

The right eigenvector is normalised such that its entries sum to one, 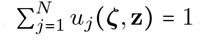. The eigenvector **u(ζ, z)** is the quasi-stationary distribution of mutant patch types as it is invariant to multiplication by **A(ζ, z)** (eq. A-3) (Harris, 1963, p. 44). Thus, *u_i_*(**ζ,z**) can be interpreted as the frequency of patches with *i* mutants among patches with at least one mutant. Left multiplying eq. (A-3) by x^T^ = (1,2,…, *N*) gives

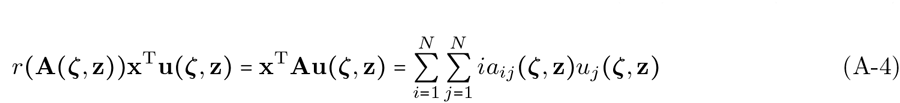

where 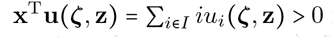 is the average number of mutants among mutant patches (i.e., patches that contain at least one mutant). Then, note that the expected total number of mutants produced over one generation by the mutants residing in a patch with *j* mutants may be written in two ways,

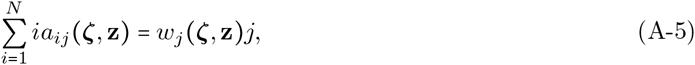

where *w_j_*(**ζ, z**) is the expected total number of adult offspring produced by a mutant over one iteration of the life cycle when there are j mutants in its patch and the remaining individuals in the population have phenotype **z**. So,

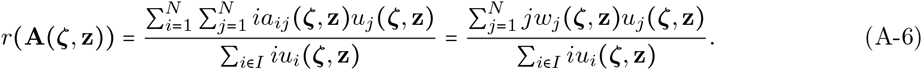

The growth rate of the mutation can therefore be written in the form of lineage fitness

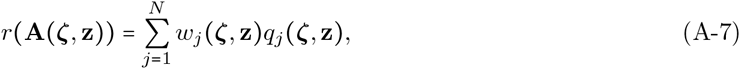

where

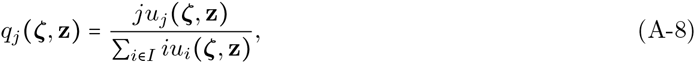

is the probability that a randomly sampled mutant from the mutant lineage belongs to a patch with *j* mutants.

#### Lineage fitness as proxy to the average growth rate and other proxies

The expression for *r*(**A(ζ, z)**) in eq. (A-7) depends on the eigenvectors of **A(ζ, z)**, which are difficult to evaluate in practice. In order to circumvent this problem, we seek a proxy whose sign around one is equivalent to that of the average growth rate, and that is also easier to evaluate. The usual fitness proxy in evolutionary biology is the basic reproductive number *R*_0_ (Ellner and Rees, 2006) which will be the starting point to our derivation.

The basic reproductive number *R*_0_ is obtained from the matrix describing mutant growth when rare, here **A(ζ, z)**, by noting that it can be decomposed as

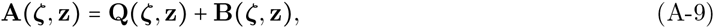

where **Q(ζ, z)** is a matrix whose element (*i,j*) gives the probability that the focal patch with *j ∊ I* mutants turns into a patch with i e I mutants. By the properties of multi-type branching processes, the transition probabilities are independent of the state of the process {**X**(*t*)}. Therefore, **Q(ζ, z)** is the transient matrix of the Markov chain describing the subpopulation of mutants in the focal patch on the state space *I*, which has the local extinction of the mutant as its only absorbing state since only residents immigrate into the patch. Meanwhile, the (*i, j*) entry of matrix **B(ζ, z)** is the expected number of patches with *i* mutants that are produced by mutant emigration from the focal patch when the latter is in state *j* (i.e., with *j* mutants).

Then, the basic reproductive number *R*_0_(**ζ, z**) is defined as the leading eigenvalue of the transition kernel giving lifetime reproduction (Caswell, 2001; Ellner and Rees, 2006), i.e., the leading eigenvalue of the matrix

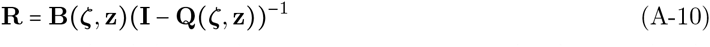

(Ellner and Rees, 2006). By construction of **A(ζ, z)** (eq. A-9) and the properties of **Q(ζ, z)** and **B(ζ, z)**, the next-generation-theorem (Thieme, 2009, Theorem 1) implies the equivalence

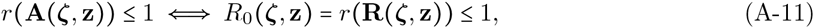

which shows that *R*_0_(**ζ, z)** is a proxy for the average growth rate, which can be computed by way of matrix inversion only.

We can further simplify the expression for *R*_0_(**ζ, z**) when only a single mutant can establish into a resident patch by immigration. In that case, only the first row of **B(ζ, z)** is non-zero, i.e.,

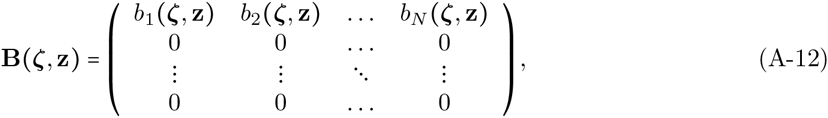

where *b_j_* (**ζ, z**) is the expected number of patches with one mutant that are produced by mutant emigration from the focal patch when in state *j*. The elements of the matrix **B(ζ, z)** below its first line are zero since the probability that two or more offspring from the same patch settle in the same patch through dispersal is zero. Note that in the infinite island model of dispersal, only a single mutant can establish into a resident patch by immigration when offspring disperse independently from one another. This can be understood by noting that if the number of patches is *N*_d_ and migration probability of mutant offspring is m, then the probability that a given breeding spot on a given patch is settled through dispersal by an offspring from the focal patch is of the order *O(m/(N_d_*)), and the probability that two or more such offspring settle in the same patch is of order *O(m^2^/(N_d_)^2^*). Summing over all patches, the probability that two or more offspring from the same individual settle on the same patch through dispersal is at most of order *O(m^2^/(N_d_)*), which goes to zero as *N*_d_ → ∞. Hence, the focal patch with *j* mutants can only turn a patch with zero mutants into a patch with a single mutant.

With **B(ζ,z)** as in eq. (A-12), the matrix **R(ζ,z)** is zero everywhere except in its first row. Since the eigenvalues of a triangular matrix are its diagonal entries, the only eigenvalue of **R(ζ, z)** that may be greater than one is simply its first diagonal element (since all the other eigenvalues are zero). In this case, the leading eigenvalue of **R(ζ, z)** (eqs. A-10-A-11) is

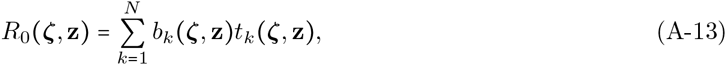

where *t*_k_(**ζ, z**) denotes the expected number of generations a patch that started with a single mutant spends with *k* mutants. Thus, when offspring disperse independently from one another, *R*_0_(**ζ, z**) is equal to the expected number of successful emigrant mutants produced by a patch that started with a single mutant, which is the definition of the proxy *R_m_*(**ζ, z**) by Metz and Gyllenberg (2001), and

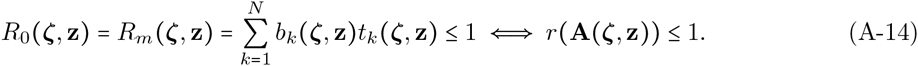

We now seek to obtain a fitness proxy expressed in terms of individual fitness, which will turn out to be of the form of lineage fitness (eq. A-7). To that end, note that the expected number of successful emigrants from a patch may be expressed as *b_k_*(**ζ,z**) = *ke_k_*(**ζ,z**), where *e_k_*(**ζ, z**) is the expected number of successful emigrants produced by a single mutant in a patch with *k* mutants when the rest of the population is monomorphic for **z**. So, we find

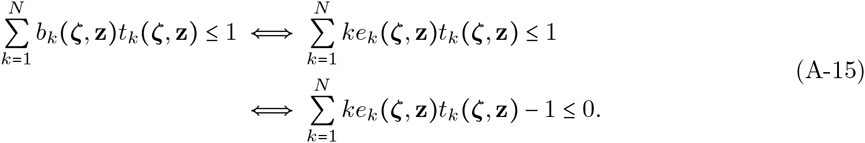

As shown in Mullon and Lehmann (2014), we can rewrite 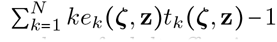 in terms of the fitness of a mutant type ζ, *w_k_* (**ζ, z**), which is the expected total number of adult offspring produced by a mutant, when there are k mutants in the patch and the remaining individuals in the population have phenotype **z**:

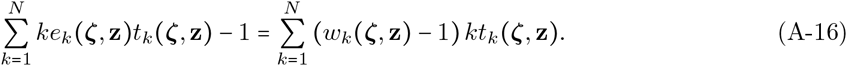

So, condition (A-15) may be expressed as

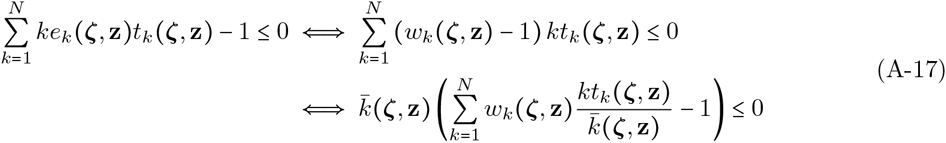

where

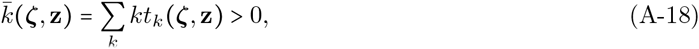

is the expected number of mutants present in a mutant patch. Now, since 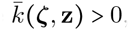, (A-17) is equivalent to

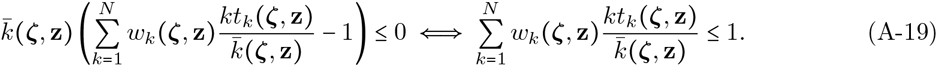

Then, note that

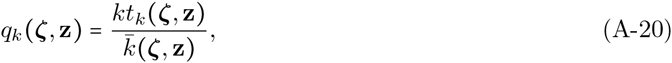

is a probability mass function which returns the probability that a randomly drawn member of the mutant local lineage resides in a patch with a total of *k ∊ {1, 2, … N}* mutants. Therefore, a mutation coding for phenotype ζ in a resident population with phenotype **z** will eventually go extinct with probability one if, and only if,

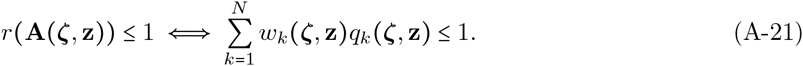

This shows that when only a single mutant can establish into a resident patch by immigration, the conditions that cause the leading eigenvalue of **A(ζ, z)** to be less or equal to one are the same as those that cause *v*(**ζ, z**) to be less or equal to one when *q_k_*(**ζ, z)** is defined in terms of sojourn times (eq. A-20) rather than the eigenvector of **A(ζ, z**) (eq. A-8). While *q_k_*(**ζ, z**) in terms of the eigenvector of **A(ζ, z)** takes into account all possible patch composition starting with different numbers of mutant immigrants, *q_k_* (**ζ, z**) in terms of sojourn times only considers a typical local lineage that starts with a single mutant. Lineage fitness *v*(**ζ, z**) with *q_k_*(**ζ, z**) given by eq. (A-20) (i.e., lineage fitness as an invasion fitness proxy) is then sufficient to characterise uninvadability.

### B. The eigenvectors of H(z) and the moulding of phenotypic correlations by selection

Here, we show that when a singular phenotype is invadable, the phenotypic correlations that are most likely to emerge are given by the eigenvector associated with the greatest positive eigenvalue of **H(z)**. Each eigenvector *e_l_*(**z**) for *l* = 1, 2, …, *n* of **H(z)** is a linear combination of the n traits. In addition, because **H(z)** is Hessian, these eigenvectors are perpendicular to one another (Horn and Johnson, 1985, p.104), i.e., *e_l_*(**z**)·e_*m*_(**z**) = 0 for *l* ≠ *m* (• refers to the dot product). They can therefore be represented as perpendicular lines in multi-dimensional phenotypic space (Supplemental Fig. 5a). The unique feature of each eigenvector *e_l_*(**z**) is that, at a singular phenotype, both the strength and direction of selection along it is only determined by its associated eigenvalue ⋋_l_(**z**). To see this, consider a mutation that appears precisely along e_1_(**z**) (Supplemental Fig. 5a). With each eigenvector scaled to have unit length 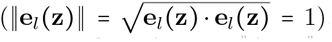, the vector **ζ − z** that connects the mutant ζ to the resident **z** can be expressed as **ζ − z** = ||**ζ - z**||e_1_(**z**). From eq. (3) and the above identities, the lineage fitness of this mutation is

**Figure 5:**
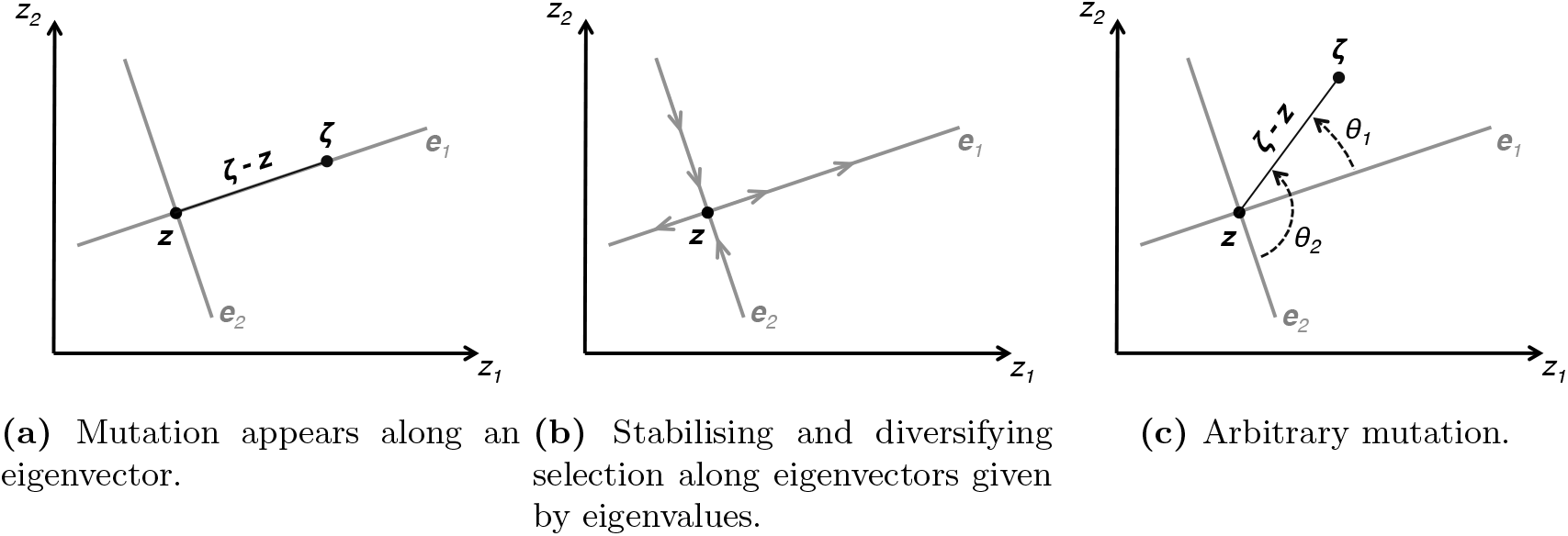
Selection close to a singular phenotype in two-traits space. The multidimensional phenotype consists of two traits, *z*_1_ and *z*_2_. The population is monomorphic for a singular phenotype **z** (black filled circle). The eigenvectors of the Hessian matrix, e_1_ and e_2_ (grey lines), are positioned to intersect at **z**. (a) A mutation appears along the eigenvector e_1_, causing the expression of trait values ζ (black filled circle). The vector that connects ζ to **z** is shown in black, (b) An example of selection direction for an invadable singular phenotype is shown. A positive eigenvalue, ⋋_1_ > 0, indicates that selection along its associated eigenvector ei is diversifying, as shown by the outward arrows. In contrast, a negative eigenvalue, *⋋_2_ <* 0, tells us that selection along e_2_ is stabilising, as shown by the inward arrows, (c) Selection on mutations away from eigenvectors. The vector that connects ζ to **z** is shown in black. The angles between this vector and both eigenvectors, *θ_1_* and *θ_2_*, are represented by dashed curved arrows.

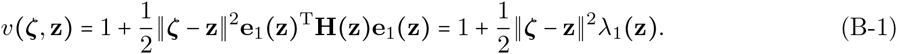

Since ||**ζ-z**||^2^ > 0, whether eq. (B-1) is greater than one only depends on the sign of the eigenvalue ⋋_1_(**z**), and in addition, the magnitude of lineage fitness is directly related to the magnitude of ⋋_1_(**z**).

The eigenvectors of **H(z)**, along with their associated eigenvalue, therefore provide a geometric representation of the direction and intensity of selection in the neighbourhood of a singular phenotype. The eigenvectors associated with negative eigenvalues give lines along which any mutation is counter-selected, whilst those associated with positive eigenvalues give lines along which selection favours mutation invasion (Supplemental Fig. 5b). In addition, the intensity of selection along an eigenvector, whether it is stabilising or diversifying, is reflected in the absolute value of its associated eigenvalue. This can be seen from the lineage fitness of a mutation that causes the expression of phenotype *ζ,* which can be written as a composite sum of the selection acting on all eigenvectors, each weighted according to the proximity of the mutation to the eigenvector,

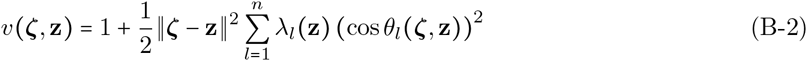

(using the eigenvalue decomposition of **H(z)**, e.g., Horn and Johnson, 1985, p. 104), where *θ_l_*(**ζ,z**) is the angle between the eigenvector of **H(z)** that is associated with *⋋_l_*(**z**), and the vector that connects the mutant phenotype ζ to the resident z in phenotypic space (i.e., cosθ_l_(**ζ, z**) = *e_l_*(**z**).(**ζ - z**)/||**ζ - z**||, see Supplemental Fig. 5c). The squared cosine of the angle θ_l_(**ζ,z**) in eq. (B-2), 0 ≤ (cos*θ_l_*(**ζ,z**))^2^ ≤ 1, measures how closely the *l*^th^ eigenvector is aligned to the vector that connects the mutant phenotype ζ to the resident **z**: when it is zero, they are perpendicular; and when it is one, they are parallel. Because all eigenvectors of a Hessian matrix are perpendicular to one another, the sum of weights is equal to one 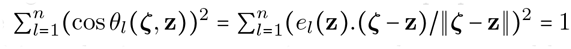. As a consequence, the maximum force that diversifying selection on a mutation can take, measured by 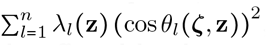, is the largest eigenvalue among the *⋋_l_*(**z**)’s, and the only mutations that will be affected by the maximum force of diversifying selection are those that appear on the eigenvector associated with the leading eigenvalue.

### C. First-order effects of selection

We derive here eq. (12) of the main text. To do so, we first reformulate the fitness function of a mutant *w_k_*(**ζ, z**), as a fitness function that depends explicitly on phenotypic values of all individuals in the population, and set 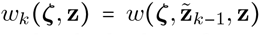 (see eq. 10). The first argument of this function is the phenotype of a focal individual whose fitness is under scrutiny, here a mutant with phenotype **ζ.** The second argument, 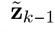, is the collection of *N* − 1 phenotypes that express the neighbours of the focal, which is composed of *k*−1 other mutants and *N−k* residents. The last argument is the resident phenotype **z** expressed in the rest of the population. Therefore, from eq. (1), we have

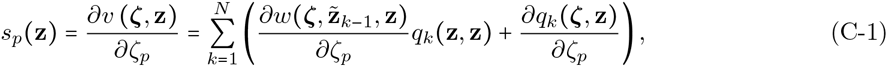

where all derivatives, here and throughout the entire appendix, are evaluated at the resident value **z**. Then, because the total probability 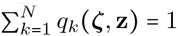 is constant, we are left with

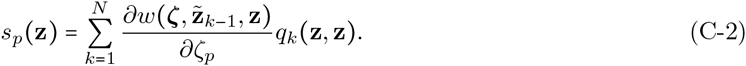

The derivative of mutant fitness with respect to mutant phenotype can be expressed in terms of individual fitness derived with respect to individual and representative neighbouring phenotypes as follows

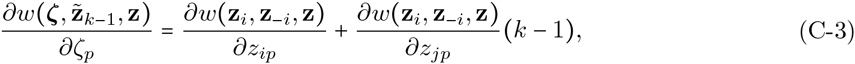

where *w*(**z_*i*_, z_*−i*_, z**) is the same function as 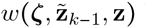 but now refers to the fitness of a focal individual, arbitrarily indexed as the individual *i ∊* {1,…, *N*}, which depends on its own phenotype z, the collection of phenotypes of its *N* − 1 neighbours **z**_*−i*_ = (**z_*i*_,…, z_*i−1*_, z_i+1_,…, z_*N*_**), as well as the resident phenotype **z** expressed in the rest of the population (e.g., Rousset, 2004). The first derivative in eq. (C-3) measures the effect of changing trait *p* in the focal *i*, and the second derivative measures the effect of changing trait *p* in a representative neighbour *j ≠ i* of the focal. Substituting eq. (C-3) into eq. (C-2), we have

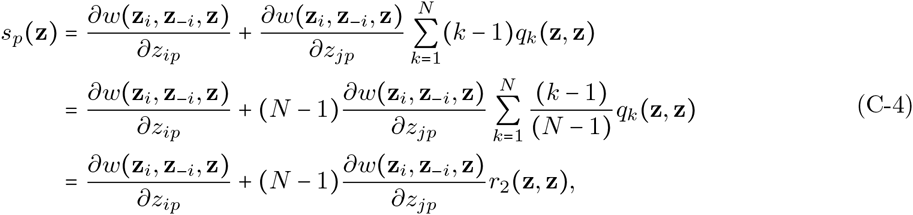

where the last line follows from the definition of *r*_2_(**z, z**) (eq. 11) and thus gives eq. (12) (see also Day, 2001, and Lehmann et al., 2015, for similar arguments).

### D. Second-order effects of selection

We derive here eq. (13) of the main text. The derivative of *v*(**ζ, z**) with respect to ζ_p_ and *ζ_q_* at ζ = **z** reads

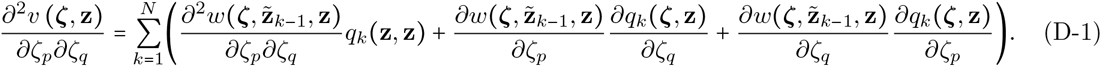

Similarly to the first derivative (eq. C-3), the second derivative of mutant fitness can be expressed in terms of individual fitness derivatives as

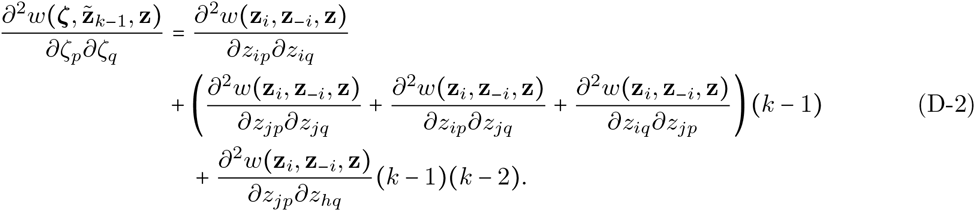

Substituting eq. (D-2) into

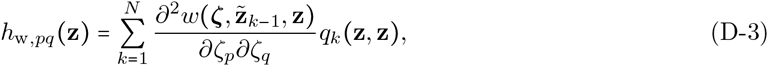

and using the definition of relatedness eq. (11), yields eq. (13b).

Substituting eq. (C-3) into eq. (D-1) and using the fact that 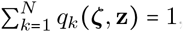, the second term of eq. (D-1) can be written as

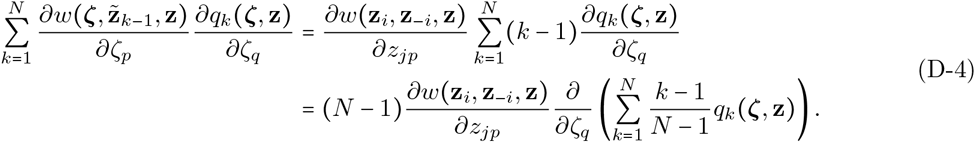

So, using the definition of *r*_2_(**ζ, z**) (eq. 11), we have

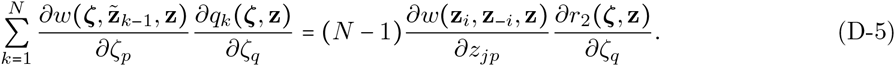

Similarly, the second term of eq. (D 1) is

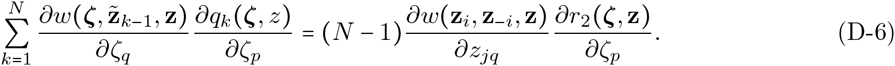

Substituting the last two equations into

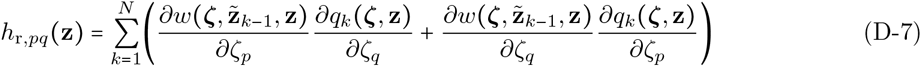

gives eq. (13c).

### E. First order effects on pairwise relatedness under a Moran process

Here, we calculate the first order effects of trait *p* on pairwise relatedness, *∂r*_2_(**ζ, z**)/∂ζ_p_ when the population follows a Moran life cycle. In this case, uninvadability can be characterised using the lineage fitness in the form of a proxy for invasion fitness. We therefore calculate *∂r*_2_(**ζ, z**)/∂ζ_p_ with the local mutant distribution *q_k_*(**ζ, z**) given by eq. (A 20). Substituting, eq. (A 20) into eq. (11), relatedness between mutants can be expressed as

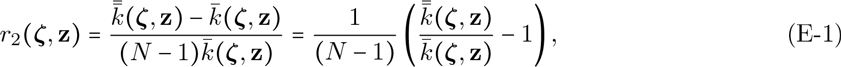

where

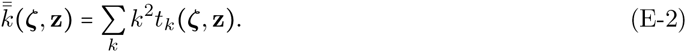

Taking the derivative of eq. (E-1) with respect to mutant effect ζ_p_ then reads

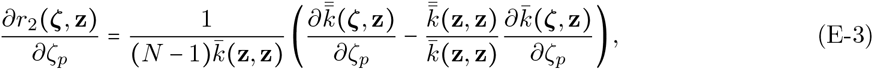

which depends on mean sojourn times under neutrality given that the patch started with a single mutant present (*t_k_* (**z, z**)) and under selection (*t_k_*(**ζ, z**)). We derive those below for the Moran process.

The general mean sojourn times *t_ij_*(**ζ, z**) spent with *i* mutants before absorption starting with *j* mutants (so that *t_k_*(**ζ,z**) = *t_k1_*(**ζ,z**)) are obtained as the elements of

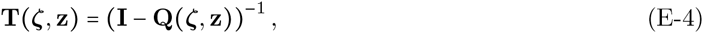

where **Q(ζ, z)** is as in eq. (A-9). For a Moran process (a birth and death process), the elements of the transient matrix **Q(ζ, z)**, i.e., the transient transition probability *q_ij_*(**ζ, z**) from *j* to *i*, can be written as

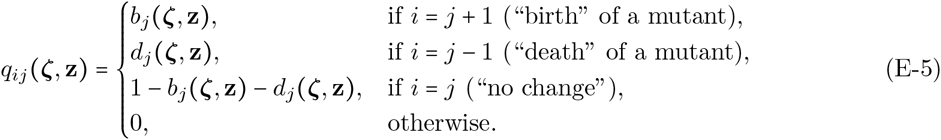

In that case, eq. (E-4) reads

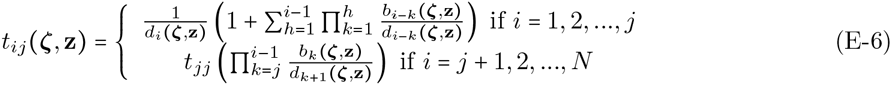

(e.g., Ewens, 2004, eq. 2.160). In particular, the mean number of generations with *i* mutants starting with a single mutant (*j = 1*) is

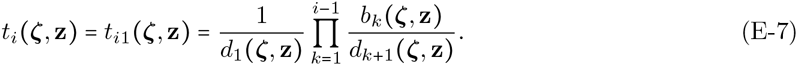

For the life cycle described in the main text (section “Uninvadability under a Moran process”), the birth and death probabilities when *k* mutants are present are

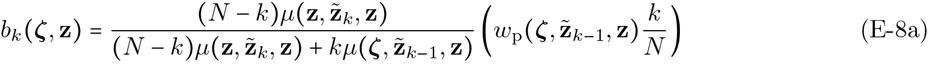

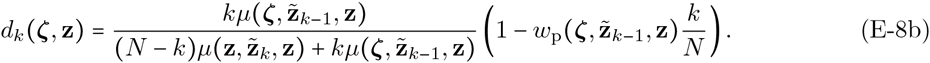

They are explained as follows and use the notation given in Box 1 in the main text. For the birth of a mutant (eq. E-8a), two events must occur. First, a resident dies, with a probability that is given by the first term of eq. (E-8a). Secondly, the offspring who settles in the vacated breeding spot descends from a mutant in that patch, which occurs with a probability that is given by the second term of eq. (E-8a). Similarly, the death of a mutant (eq. E-8b) requires the death of a mutant and replacement by a resident.

The neutral sojourn times *t_k_* (**z, z**) are found using eq. (E-8) substituted into eq. (E-7) evaluated at **ζ = z**. Doing so, we find

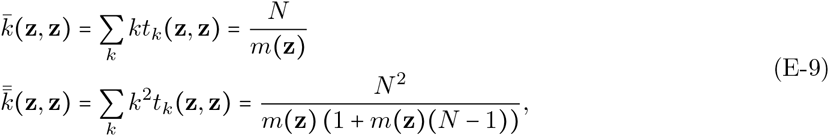

where *m*(**z**) = 1 - *w*p (**z z z**) is the backward migration rate.

The first order effects of trait *p* on pairwise relatedness (eq. E-3) also depends on the sojourn times under selection, 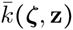 (eq. A-18) and 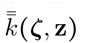 (eq. E-2). These terms can be rewritten as the following matrix operation,

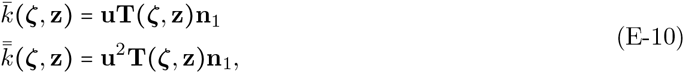

where **u** = (1, 2, …, *N*), **u**^2^ = (1,4,…, *N*^2^) and **n**_*1*_ = (1,0,…, 0)^T^. The first order effect of a trait on relatedness (eq. E-3) depends on the derivatives of eq. (E-10) with respect to mutant effect. Using eq. (E-4), and the fact that the derivative of the inverse of a matrix A is given by

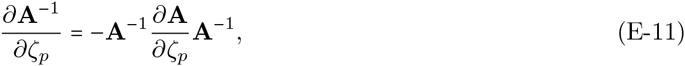

the derivatives of eq. (E-10) with respect to mutant effect *ζ_p_* are

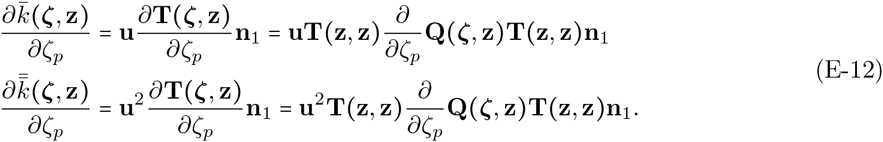

Now, using eqs. (E-6)-(E-8) at neutrality (**ζ = z**), some algebraic manipulations show that

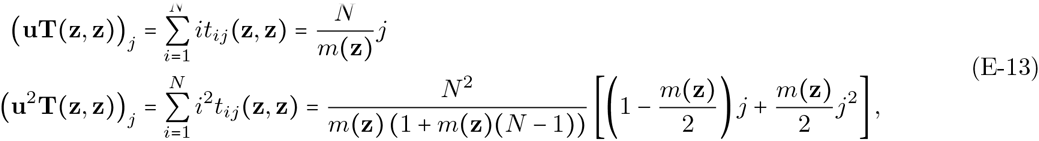

which in matrix form reads

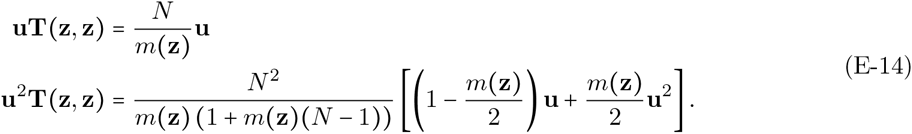

Substituting eq. (E-14) into eq. (E-12) then gives

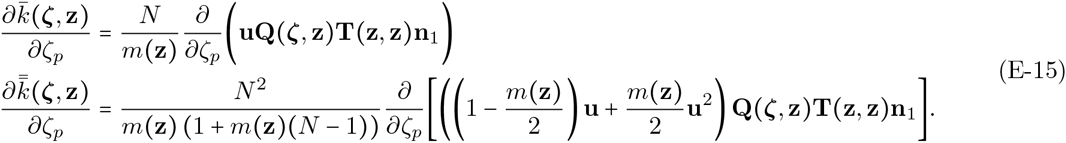

Writing *ϕ*_1,*j*_ (**ζ, z**) and *ϕ*_2,*j*_ (**ζ, z**) as the first and second moments of the number of mutants in the focal patch after a generation conditional on there being j mutants before the transition, we have

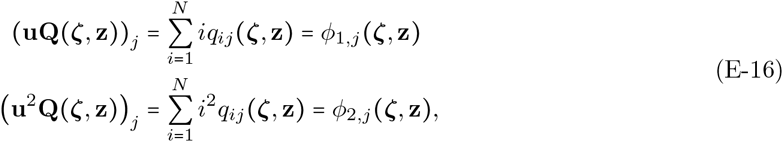

and by definition,

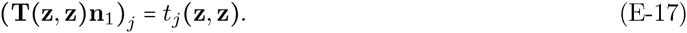

Thus, eqs. (E-16)-(E-17) substituted into eq. (E-15) give

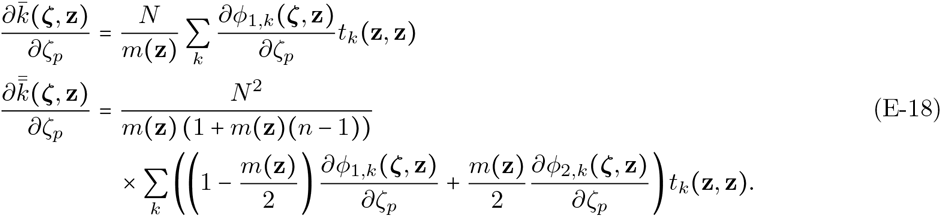

We now find an expression for the derivatives of *ϕ_1,k_* (**ζ, z**) and *ϕ_2,k_*(**ζ, z**). Conditional on *k*, the first and second moments of the number of mutants in the focal patch after a life cycle iteration are

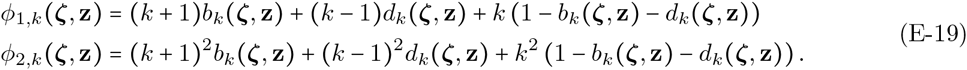

In order to get the derivatives of eq. (E-19) with respect to *ζ_p_*, we substitute eq. (E-8), use the chain rule and the following identities,

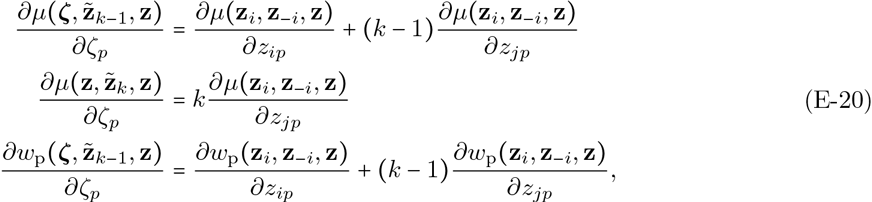

and eventually obtain

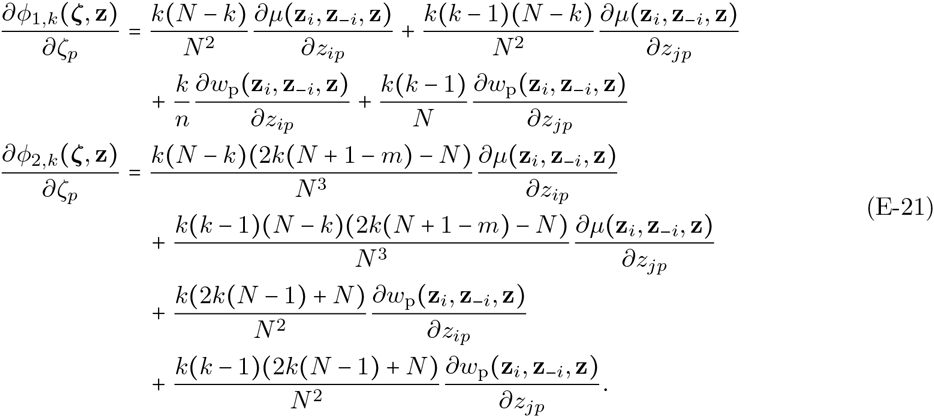

Finally, we substitute eq. (E-21) into eq. (E-18), which is in turn substituted into eq. (E-3), and we use the first and second moments of sojourn times (eq. E-9), as well as the third and fourth moments,

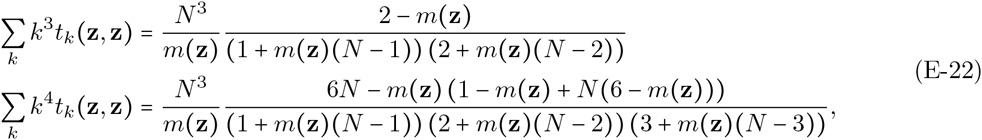

to obtain the result of eq. (14) and Table 2 of the main text.

The first order effects of a trait on pairwise relatedness that we have derived for the Moran process (eq. 14) can be written in a form similar to the one that has been used for the Wright Fisher process (Ajar, 2003; Wakano and Lehmann, 2014). In the absence of variation among death rates, *μ*(**z_*i*_, z_*−i*_, z**) = *μ* for all *i*, eq. (14) with Table 2 can be expressed as

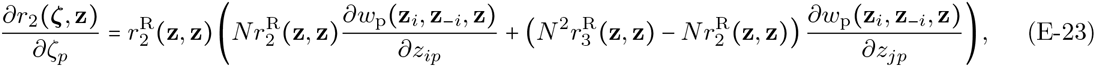

where

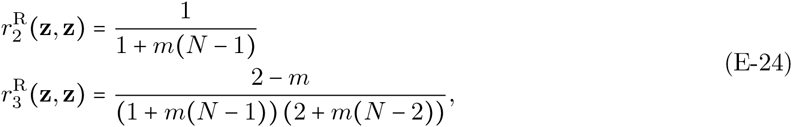

are the probabilities of sampling two and three individuals with replacement that are identical-by-descent respectively. Eq. (E-23) bears close to resemblance to the parts of eq. (18) of Ajar (2003) and eq. (C.2) of Wakano and Lehmann (2014) that correspond to the first order effects of a trait on pairwise relatedness for the Wright Fisher process. Differences with eq. (E-23) are due to the fact that generations overlap in the Moran process, but not in the Wright Fisher process.

We now show that the first order effects of a trait on pairwise relatedness vanishes at singular phenotypes if the evolving traits only affect adult fertility or offspring survival. If that is the case, then the death rate *μ*(**z_*i*_, z_*−i*_, z**) = *μ* constant for all individuals in the population, and the philopatric component of fitness may be written as eq. (15). Substituting eq. (15) into eq. (14) and using Table 2, we find that

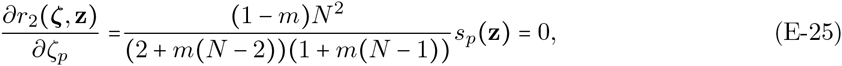

since at a singular phenotype, *s_p_*(**z**) = 0.

### F. Neutral pairwise and three-way relatedness for a Moran process

The stability criteria also depend on the probabilities of that two and three individuals are related in the resident population. These are found using standard identity-by-descent arguments (e.g., Karlin, 1968), by solving the recursions

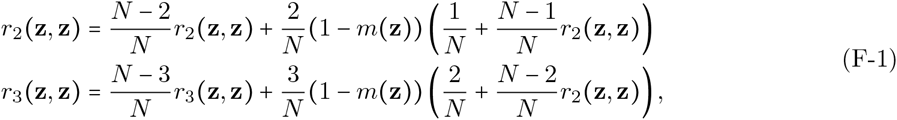

for a Moran process, whose solutions are given in Table 1.

### G. Uninvadability under weak effects

In order to derive the selection gradient and Hessian matrix when payoffs have weak effects (eqs. 18 … 19), we follow the line of arguments developed in Lehmann et al., 2015 and first observe that under weak effects (i.e., *∊* → 0 in eq. 17), the fitness of a focal mutant in the focal patch with *k* mutants can be expressed as

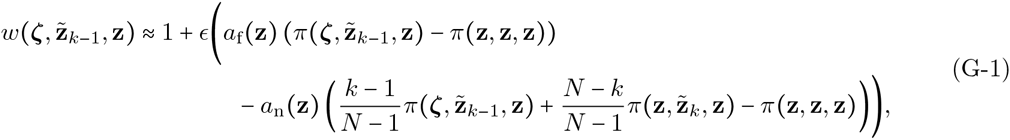

to the first order of *∊* (and where we used the zero-sum effect of selection on fitness in populations of constant size, see p. 96 of Rousset, 2004, for explanation). The difference 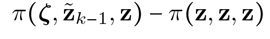 is the diference between the payof received by the focal and the payof of a resident from another patch, and 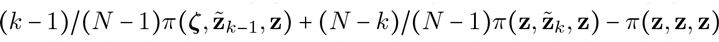 is the difference between the average payoff in the focal patch, excluding the focal individual, and the payof of a resident from another patch. So, the diference between these two diferences measures how well the focal does compared to the rest of her patch, relative to an individual from another patch. Focal fitness depends on the diference between these two, but where each is weighted by a coefficient, a_f_(**z**) and a_n_(**z**) respectively, that together measure the spatial scale of competition in the resident population (Frank, 1998).

The spatial scale of competition is given by 0 ≤ a(**z**) = a_n_(**z**)/a_f_(**z**) ≤ 1 (since 0 < a_f_(**z**) ≤ 1 and a_n_(**z**) ≤ a_f_(**z**) according to our assumptions listed in the main text, see “Uninvadability under weak selection” section). To see how a(**z**) measures the spatial scale of competition, we rewrite eq. (G-1) as

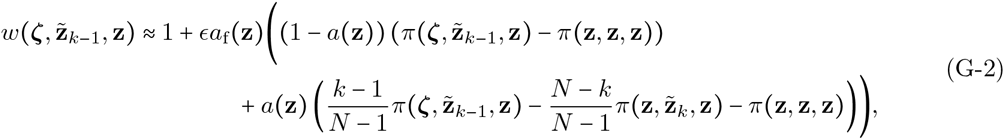

which shows that if a(**z**) is small, then the payof of neighbours has little efect on focal fitness compared to the payof of individuals from other patches, i.e., competition tends to occur globally. This would occur, for instance, if dispersal is strong. Conversely, if a(**z**) is close to one, then competition tends to occur locally. The coefficients a_f_(**z**) and a_n_(**z**), and therefore a(**z**), are model specific, and will depend on the demographic properties of the population, such as patch size, dispersal rate, or the number of open breeding spots at each generation, as well as the evolving trait. The coefficients a_f_(**z**) and a_n_(**z**) are found by Taylor expanding the fitness function in the form of eq. (G-1).

Lineage fitness (eq. 1) is found by marginalising eq. (G-1) over the distribution of mutants *q_k_*(**ζ,z**). However, since e is small and eq. (G-1) is of *O(∊)*, it is sufficient to marginalise over the probability mass function for the number of mutants under neutrality, 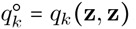,

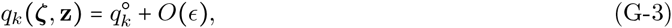

where 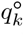 is independent of the mutant type. Substituting eq. (G-1) and eq. (G-3) into lineage fitness eq. (1) gives to the order of *O(∊)*

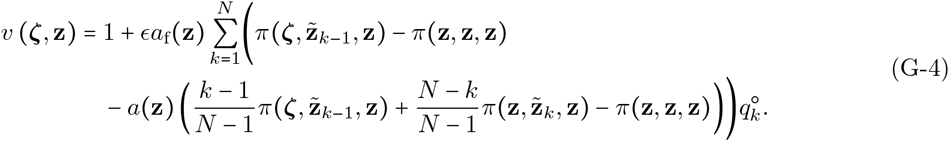

*First-order condition*. Taking the derivative of eq. (G-4) with respect to *ζ_p_* at ζ = **z** reads

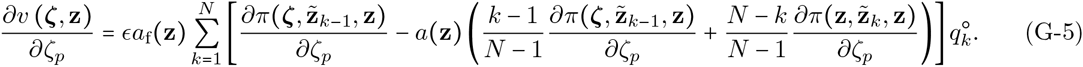

Then, after substituting for the derivatives,

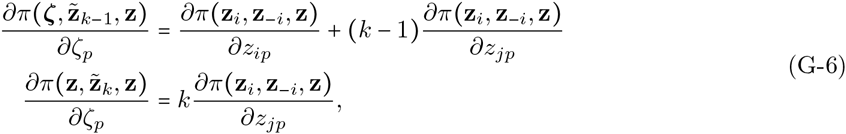

some re-arrangement, and using the definition of relatedness eq. (11), eq. (G-5) gives the result eq. (18). *Second-order condition*. Taking the derivative of eq. (G-4) with respect to *ζ_p_* and *ζ_q_* at ζ = **z** yields

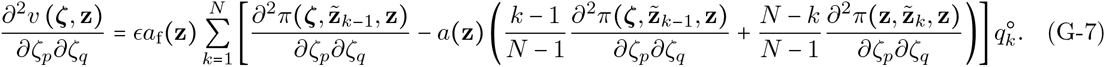

The derivatives can be expressed as

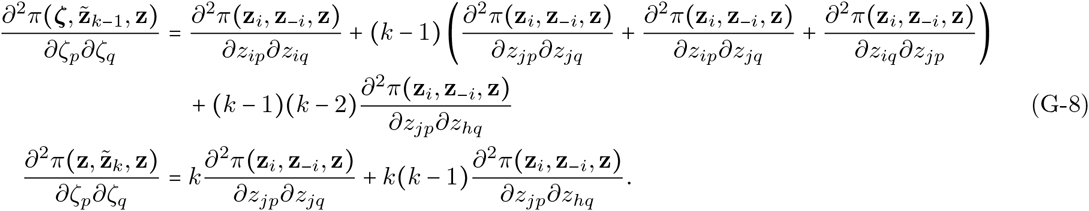

Substituting eq. (G-8) into eq. (G-7) and using the definitions of relatedness eqs. (11) and Table 1 generates the result eq. (19).

### H. Agreement between weak and strong selection modes

Here, we show that the sign of *h*_11_(**z**) derived under weak selection (eq. 23) is the same as the sign of *h*_11_(**z**) for strong selection (equation not shown). In order to do so, we used Mathematica 10.0.1.0 (Wolfram Research, 2014) to generate random values for the parameters *b*_1_, *b*_2_, *c*_2_ and *c*_2_ between -100 and 100, for the patch size *N* between 2 and 1000 and for effective dispersal m between 0 and 1 (code available on request). Then, if these values generated admissible singular helping strategies eq. 22, (i.e., between 0 and 1), and admissible individual fertility at singular helping strategy eq. 21, (i.e., positive), then we compared the sign of *h*_11_(**z**) derived under the weak and strong selection modes. Out of 2.10^6^ random sample, 465943 produced admissible singular strategies and individual fertility and, in all cases, the sign of *h*_11_(**z**) between weak and strong selection modes matched.

We show similarly that the sign of h_11_ given by eq. 23 and the sign of the leading eigenvalue of **H(z)** with e = 1 (equation not shown) are very often equivalent. Out of 2.10^6^ random sample, 465359 produced admissible singular strategies and individual fertility and, in 464686 cases (99.86%), the sign of h_12_(z) given by eq. (32) matched the sign of h_12_(z) for strong selection and arbitrary patch size.

We also show that the sign of *h*_12_(**z**) derived under weak selection and large patch size (eq. 32) is very often the same as the sign of *h*_12_(**z**) for strong selection and arbitrary patch size (equation not shown). Out of 2. 10^6^ random sample, 465600 produced admissible singular strategies and individual fertility and, in 465589 cases (99.998%), the sign of *h*_12_(**z**) given by eq. (32) matched the sign of *h*_12_(**z**) for strong selection and arbitrary patch size. We remark here that *h*_12_(**z**) to the first order of selection *∊* but with arbitrary patch size is

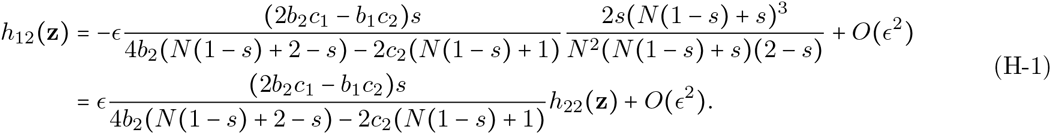

### I. Individually-based simulations

We used Mathematica 10.0.1.0 (Wolfram Research, 2014) to simulate the joint evolution of helping and dispersal in a population with *N_p_* = 1000 patches, each populated by *N* = 8 individuals (code available on request). Starting with a population monomorphic for singular helping and dispersal strategies, we track the evolution of the multidimensional phenotypic distribution as small mutations continuously arise. Each individual is characterised by a level of helping *x*_*i*_ and dispersal probability *d*_*i*_. At the beginning of a generation, we calculate the fertility *f*_*i*_ of each individual according to its helping strategy and that of its neighbours (eq. 21 with e = 1). We use parameter values for the benefit and cost of helping (*b*_1_ = 6, *b*_2_ = −1.4, *c*_1_ = 4.56, *c*_2_ = −1.6) that are known to lead to evolutionary branching in infinite well-mixed populations (Doebeli et al., 2004). Then, in each patch, an individual is chosen at random to die. We replace it by means of the weighted sampling of an individual in the population, where each individual is weighted according to whether they belong to the patch on which the breeding spot is filled or not. If an individual belongs to the same patch in which a breeding spot is filled, then its weight is *f_i_*(1−*d*_i_). If it belongs to another patch, then its weight is *f*_*i*_*d*_*i*_/(*N*_p_ − 1). Once an individual is chosen to fill the breeding spot, its phenotypic values mutate with probability 0.01. If they do not mutate, then the offspring has the same phenotypic values as its parents. If they mutate, then we add small perturbations to the parental level of helping and dispersal probability that are sampled from a binormal distribution with mean ( 0, 0), variance 0. 02^2^ in each trait, and no covariance. The resulting phenotypic values are controlled to remain between 0 and 1. Once a breeding spot has been opened and filled in each patch, the generation is over and we repeat the iteration. For each values of *s ∊* {0.04,0.2,0.5,0.8,0.91,0.92,0.93,0.94,0.95,0.96}, the population is initially monomorphic for the singular strategies and we simulate 1.5 x 10^5^ generations.

To simulate the evolution of helping alone, the same procedure as above is used, except that survival in dispersal is fixed at *s* = 1 and dispersal is held constant at *m ∊* {0.6,0.68,0.69,0.71} for each individual. In the case of a mutation, only helping is perturbed by an amount sampled from a Normal distribution with mean 0 and variance 0. 02^2^.

### J. First order perturbation of the eigenvectors and eigenvalues of a matrix

Here, we derive eqs. 30 and 31 from the main text. To do this, we note that if we can express the Hessian matrix as **H(z) = H^*^(z) + *∊*H^•^(z)** where *∊* is small, and we label 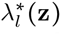 and 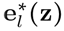, the *l*^th^ eigenvalue of **H*(z)** and its associated eigenvector respectively, then the corresponding eigenvalues and eigenvectors of **H(z)** are approximately

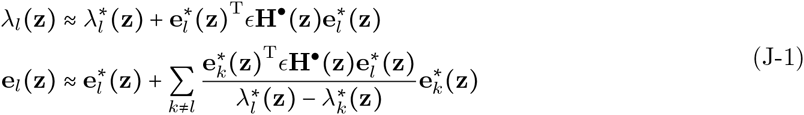

(Golub and Van Loan, 1996, p. 323). In our example of the joint evolution of helping and dispersal, we find that substituting fitness (eq. I-a, Box 1) into eq. (13) together with the perturbation of relatedness eq. (14) and evaluating it at eq. (29), to the first order of e the Hessian matrix can be decomposed as **H(z) = H^*^(z) + *∊*H^•^(z)** where

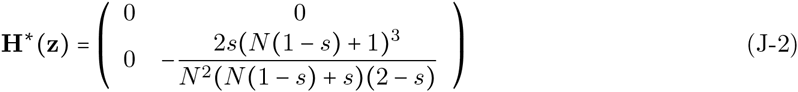

and

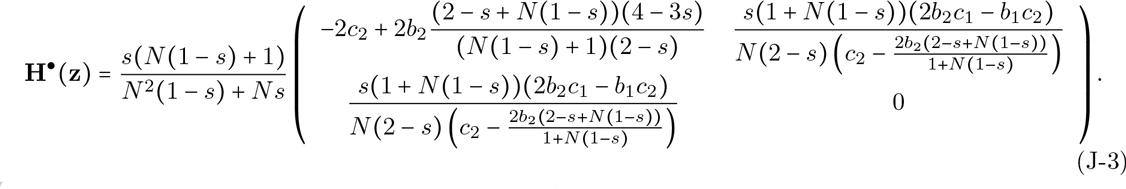

The eigenvalues and eigenvectors of **H^*^(z)** are then

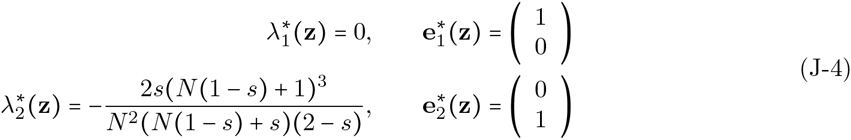

which substituted into eq. (J-1) give the approximate results of the main text (eqs. 30 and 31).

